# *Ustilago maydis* serves as a novel production host for the synthesis of plant and fungal sesquiterpenoids

**DOI:** 10.1101/2020.05.04.076794

**Authors:** Jungho Lee, Fabienne Hilgers, Anita Loeschke, Karl-Erich Jaeger, Michael Feldbrügge

## Abstract

Sesquiterpenoids are important secondary metabolites with various pharma- and nutraceutical properties. In particular, higher basidiomycetes possess a versatile biosynthetic repertoire for these bioactive compounds. To date, only a few microbial production systems for fungal sesquiterpenoids have been established. Here, we introduce *Ustilago maydis* as a novel production host. This model fungus is a close relative of higher basidiomycetes. It offers the advantage of metabolic compatibility and potential tolerance for substances toxic to other microorganisms. We successfully implemented a heterologous pathway to produce the carotenoid lycopene that served as a straightforward read-out for precursor pathway engineering. Overexpressing genes encoding enzymes of the mevalonate pathway resulted in increased lycopene levels. Verifying the subcellular localisation of the relevant enzymes revealed that initial metabolic reactions might take place in peroxisomes: despite the absence of a canonical peroxisomal targeting sequence, acetyl-CoA C-acetyltransferase Aat1 localised to peroxisomes. By expressing the plant (+)-valencene synthase CnVS and the basidiomycete sesquiterpenoid synthase Cop6, we succeeded in producing (+)-valencene and α-cuprenene, respectively. Importantly, the fungal compound yielded about tenfold higher titres in comparison to the plant substance. This proof of principle demonstrates that *U. maydis* can serve as promising novel chassis for the production of terpenoids.

## Introduction

Terpenoids (isoprenoids) constitute an important class of secondary metabolites found mainly in plants and fungi. They are classified according to the number of C5 scaffold isopentenyl diphosphate (IPP) building blocks. Sesquiterpenoids, for example, contain three and diterpenoids consist of four of such building blocks, forming C15 and C20 scaffolds, respectively. Terpenoids exhibit a plethora of biological functions like photoprotection, hormone signalling and defence against pathogens (Schmidt-Dannert, 2015;Troost et al., 2019;Moser and Pichler, 2019). Because of these diverse bioactivities they are also of interest for biotechnology. Artemisin, for example, functions as antimalarial drug and (-)-patchoulol serves as a valuable fragrance for the cosmetics industry. Lycopene and (+)-valencene are natural food additives (Schempp et al., 2018;Moser and Pichler, 2019).

Traditionally, terpenoids are extracted from plant or fungal materials. Extensive efforts are made to produce these compounds in heterologous microorganisms for increased sustainability as well as to expand the chemical diversity of terpenoids by further modification of near-to-natural versions (Ro et al., 2006;Ignea et al., 2018;Schempp et al., 2018). Several companies like Amyris, Evolva, Isobionics (now BASF) and Firmenich have already marketed diverse terpenoids produced with engineered microorganisms (Schempp et al., 2018).

A rich source of terpenoids is higher basidiomycetes, forming mushrooms (Schmidt-Dannert, 2015;Xiao and Zhong, 2016). Prominent examples are derivatives of the sesquiterpenoid illudin S from *Omiphalotus olearius* that exhibits strong anti-tumour activity (Jaspers et al., 2002) and the diterpenoid pleuromutilin from *Clitopilus passeckerianus* that has an antibacterial effect by inhibiting the large subunit of prokaryotic ribosomes (Hartley et al., 2009).

Most fungal terpenoids are not yet studied because it is difficult to obtain enough pure substances for defined biological assays. For the current access of the compounds, researchers rely mostly on improving the biosynthesis in the natural producer. However, higher basidiomycetes often entail the disadvantage of being difficult to cultivate and that sophisticated molecular tools for pathway engineering are not available. Alternatively, synthetic hosts like *Escherichia coli* or *Saccharomyces cerevisiae* are used (Xiao and Zhong, 2016). Despite their advantages in biotechnology, the production of antibacterial substances in *E. coli* is challenging and terpenoids from basidiomycetes might be toxic for ascomycetes like *S. cerevisiae*. Based on these reasons, we followed the strategy to exploit a well-studied basidiomycete for the production of terpenoids. We chose the corn smut *Ustilago maydis* that serves as an excellent model system for basic cell biology and plant pathology (Kahmann and Kämper, 2004;Zarnack and Feldbrügge, 2010;Béthune et al., 2019). The genome is sequenced and well-annotated. Transcriptomics and proteomics have been carried out and sophisticated molecular tools are established (Kämper et al., 2006;Scherer et al., 2006;Koepke et al., 2011;Olgeiser et al., 2019). Strains can be efficiently generated by homologous recombination and after stable insertion in the genome, selection markers can be excised resulting in marker-free strains either to recycle the resistance marker or to obtain safe strains for biotechnological production (Khrunyk et al., 2010;Terfrüchte et al., 2014).

Besides serving as a model for basic research, *U. maydis* is also advancing as a flexible microorganism for biotechnological applications. Importantly, the yeast form is non-pathogenic. Furthermore, *U. maydis* only infects corn and infected ears have been eaten as a delicacy for centuries in Mexico, indicating that its consumption is not harmful to humans (Feldbrügge et al., 2013). The fungus constitutes a promising production chassis for a whole variety of biotechnologically relevant compounds including itaconic acid, a versatile building block for tailor-made biofuels (Becker et al., 2019;Wierckx et al., 2019). Additionally, it produces biosurfactants like mannosylerythritol lipids and ustilagic acid, which can be used as basis for sustainable detergents and emulsifiers (Teichmann et al., 2007;Teichmann et al., 2010;Feldbrügge et al., 2013). A recent application is the establishment of *U. maydis* as a novel host for the expression of heterologous proteins. This is strongly supported by the discovery that valuable proteins like antibody formats can be exported in the culture medium by a novel unconventional secretion pathway (Sarkari et al., 2014). This pathway prevents undesired glycosylation of the product that is usually processed during conventional secretion (Stock et al., 2012;Sarkari et al., 2014;Terfrüchte et al., 2018).

At present, *U. maydis* strains are being optimised to convert plant biomass into valuable products. Strains have been successfully engineered to grow on cellulose, xylose and polygalacturonic acid (Geiser et al., 2016a;Stoffels et al., 2020). The latter is a major component of pectin. Thus, a consolidated bioprocess is being developed, in which complex natural substrates are converted to fermentable sugars and to value-added compounds in the future (Geiser et al., 2016a). Here, we add terpenoids to the growing list of compounds produced in *U. maydis*.

## Results

### *U. maydis* contains an evolutionarily conserved FPP pathway

To design a strategy for the heterologous production of sesquiterpenoids in *U. maydis*, we predicted the underlying metabolic pathways using bioinformatics analysis (Figure 1A, Supplementary Figure S1). As a starting point, we adopted information from the KEGG pathway “terpenoid backbone biosynthesis” for *U. maydis* (Kyoto Encyclopedia of Genes and Genomes, https://www.genome.jp/kegg/). We conceptually divided the metabolic network into four parts: the mevalonate module, the prenyl phosphate module, the carotenoid module and the recombinant sesquiterpenoid module (Troost et al., 2019). The mevalonate pathway from acetyl-CoA to isopentyl- and dimethylallyl diphosphate (IPP and DMAPP) is evolutionarily highly conserved in eukaryotes (Miziorko, 2011). Therefore, we used the detailed knowledge on *S. cerevisiae*, *H. sapiens*, and *A. thaliana* as a blueprint (Figure 1A, Table 1; Supplementary Figure S1; Nielsen and Keasling, 2016;Ye et al., 2016). The enzymatic functions of enzymes Hcs1 (3-hydroxy-3-methylglutaryl-CoA-synthase) and Hmg1 (3-hydroxy-3-methylglutaryl-coenzyme A reductase) have previously been studied in *U. maydis* (Croxen et al., 1994;Winterberg et al., 2010). The remaining enzymes (acetyl-CoA-C-acetyltransferase Aat1, mevalonate kinase Mvk1, phosphomevalonate kinase Pmk1, mevalonate diphosphate decarboxylase Mdc1, isopentenyl diphosphate isomerase Idi1and farnesyl pyrophosphate synthase Fps1) were identified by high amino acid sequence similarity and the presence of conserved domains when compared to well-studied fungal, human and plant versions (Figure 1A, Table 1; Supplementary Figure S1). Mevalonate kinase was the only enzyme not predicted in the KEGG pathway database. The same strategy was applied for the identification of enzymes of the prenyl phosphate module, producing geranylgeranyl-diphosphate via chain elongation from IPP and DMAPP (GGPP; Figure 1A, Table 1).

**Table 1.** Enzymes of the mevalonate pathway in different organisms

| *U. maydis* |  |  | *S. cerevisiae* |  |  | *H. sapiens* |  |  |
| --- | --- | --- | --- | --- | --- | --- | --- | --- |
| Name | UMAG | NCBI annotation | Name | Identities (%) | e-value | Name | Identities (%) | e-value |
| Aat1 | 03571 | Acetyl-CoA C-acetyltransferase | Erg10p | 209 (52) | 2e-128 | ACAT1 | 240 (59) | 4e-166 |
|  |  |  |  |  |  | ACAT2 | 175 (45) | 7e-104 |
| Hcs1 | 05362 | Probable hydroxymethylglutaryl-CoA synthase | Erg13p | 219 (48) | 1e-154 | HMGCS1 | 234 (50) | 3e-154 |
|  |  |  |  |  |  | HMGCS2 | 228 (49) | 1e-147 |
| Hmg1 | 15002 | Probable 3-hydroxy-3-methylglutaryl-coenzyme A reductase | Hmg1p | 273 (62) | 1e-179 | HMGCR | 258 (60) | 8e-172 |
|  |  |  | Hmg2p | 278 (60) | 0 |  |  |  |
| Mvk1 | 05496 | Mevalonate-5-kinase | Erg12p | 147 (37) | 1e-60 | MVK | 148 (38) | 3e-53 |
| Pmk1 | 00760 | Phosphomevalonate kinase | Erg8p | 150 (29) | 2e-40 | PMVK | No homologs found |  |
| Mdc1 | 05179 | Mevalonate-5-pyrophosphate decarboxylase | Mvd1p | 198 (50) | 2e-119 | MVD | 200 (49) | 3e-112 |
| Idi1 | 04838 | Isopentenyl diphosphate isomerase | Idi1p | 137 (52) | 2e-77 | IDI1 | 116 (53) | 3e-68 |
|  |  |  |  |  |  | IDI2 | 98 (42) | 2e-51 |
| Fps1 | 11875 | Putative bifunctional (2E,6E)-farnesyl diphosphate synthase/dimethylallyl transtransferase | Erg20p | 211 (69) | 9e-155 | FDPS | 169 (50) | 8e-103 |

**Figure 1.**
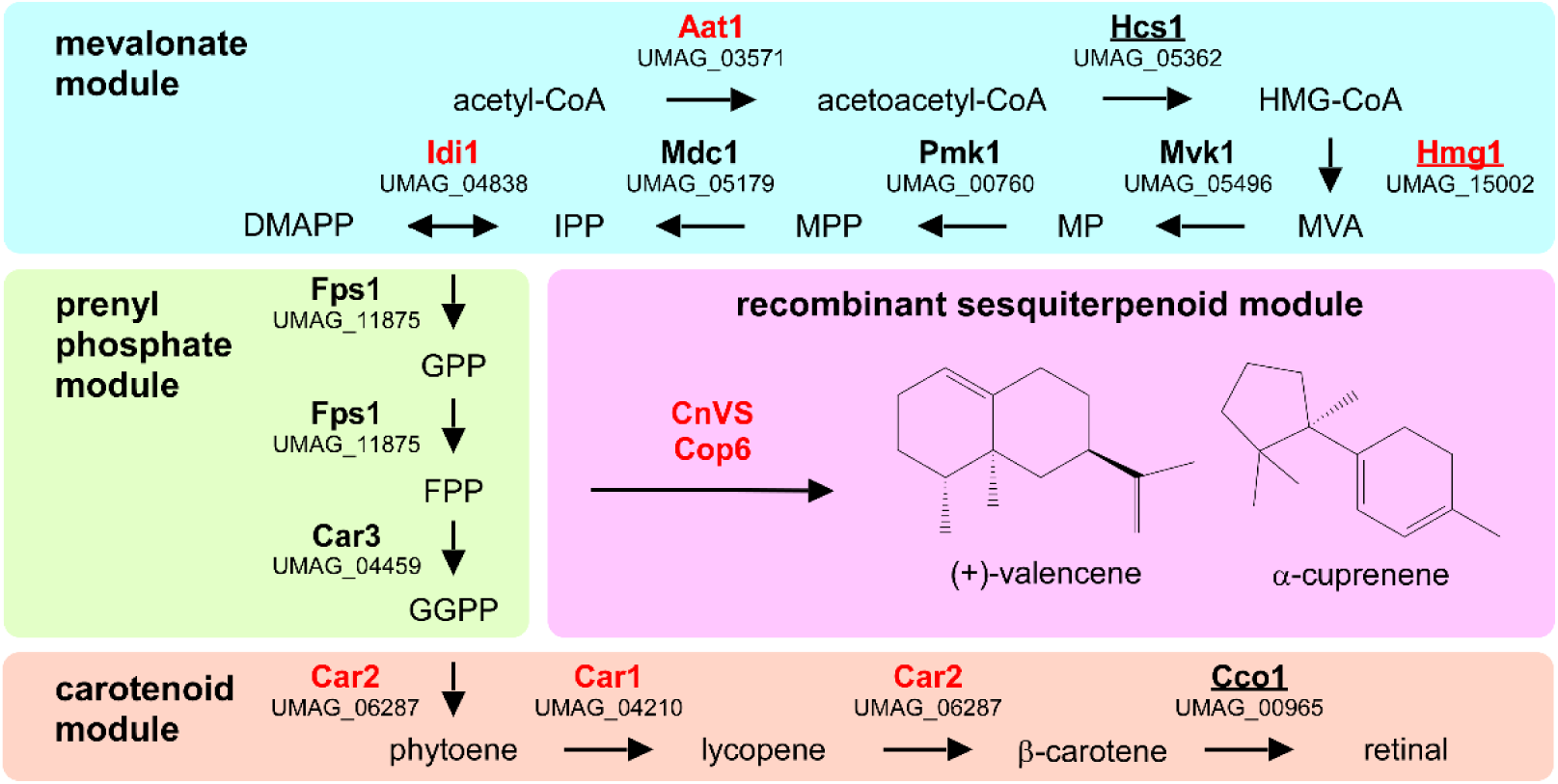
Metabolic network for the heterologous production of sesquiterpenoids. Graphical representation of the various modules involved in recombinant sesquiterpenoid synthesis: mevalonate module (blue), prenyl phosphate module (green), carotenoid module (orange) and recombinant sesquiterpenoid module (pink). Enzyme names are given in Table 1. Enzymes studied in this publication are indicated in red and those that were already studied in U. maydis are underlined (HMG-CoA, 3-hydroxy-3-methylglutrayl-CoA; MVA, mevalonate; MP, mevalonate-5-phosphate; MPP, mevalonate-pyrophosphate; IPP, isoprenylpyrophosphate; DMAPP, dimethylallyl-pyrophosphate, GPP, geranyl-pyrophosphate; FPP, farnesyl-pyrophosphate GGPP, geranylgeranylpyrophosphate).

The carotenoid module has been predicted before and its main function is the production of retinal that serves as chromophore for photoactive opsin channels Ops1-3 (Estrada et al., 2009;Panzer et al., 2019). The cleavage reaction of β-carotene into two retinal molecules is catalysed by Cco1 (β-carotene cleavage oxygenase; Estrada et al., 2009). It was already shown that deletion of *cco1* resulted in the accumulation of β-carotene (Estrada et al., 2009). Loss of retinal synthesis causes no mutant phenotype under standard growth conditions or during pathogenic development, thus the biological function of opsins in *U. maydis* is as yet unclear (Estrada et al., 2009;Panzer et al., 2019).

For the recombinant sesquiterpenoid module, we aimed to branch off the key precursor farnesylpyrophosphate (FPP). We chose the plant (+)-valencene synthase from *Callitropsis nootkatensis* (CnVS; Troost et al., 2019) and the fungal α-cuprenene synthase Cop6 from *Coprinopsis cinerea* (Agger et al., 2009) to synthesise (+)-valencene and α-cuprenene, respectively (Figure 1A). In essence, (+)-valencene served as a benchmarking product for our new approach as it has been the target in multiple studies on microbial sesquiterpenoid production before, while α-cuprenene served as an example of a basidiomycete compound that has been well-studied before (Agger et al., 2009;Beekwilder et al., 2014;Frohwitter et al., 2014;Stöckli et al., 2019;Troost et al., 2019).

### Establishing the production of lycopene in *U. maydis* as an indicator of carotenoid precursors

For the production of recombinant sesquiterpenoids, we aimed to increase the activities of the mevalonate module in order to obtain higher levels of the key precursor FPP (Figure 1A). However, FPP is a toxic intermediate and the detection of intracellular FPP levels is not trivial (Dahl et al., 2013). Therefore, we decided to modify the intrinsic carotenoid module to accumulate lycopene, a coloured FPP-derived product that can be easily detected and quantified (Figure 2A-B). Carotenoid synthesis should serve as an easy read-out system for the activity of the underlying metabolic pathway as well as a safety valve for high FPP levels.

**Figure 2.**
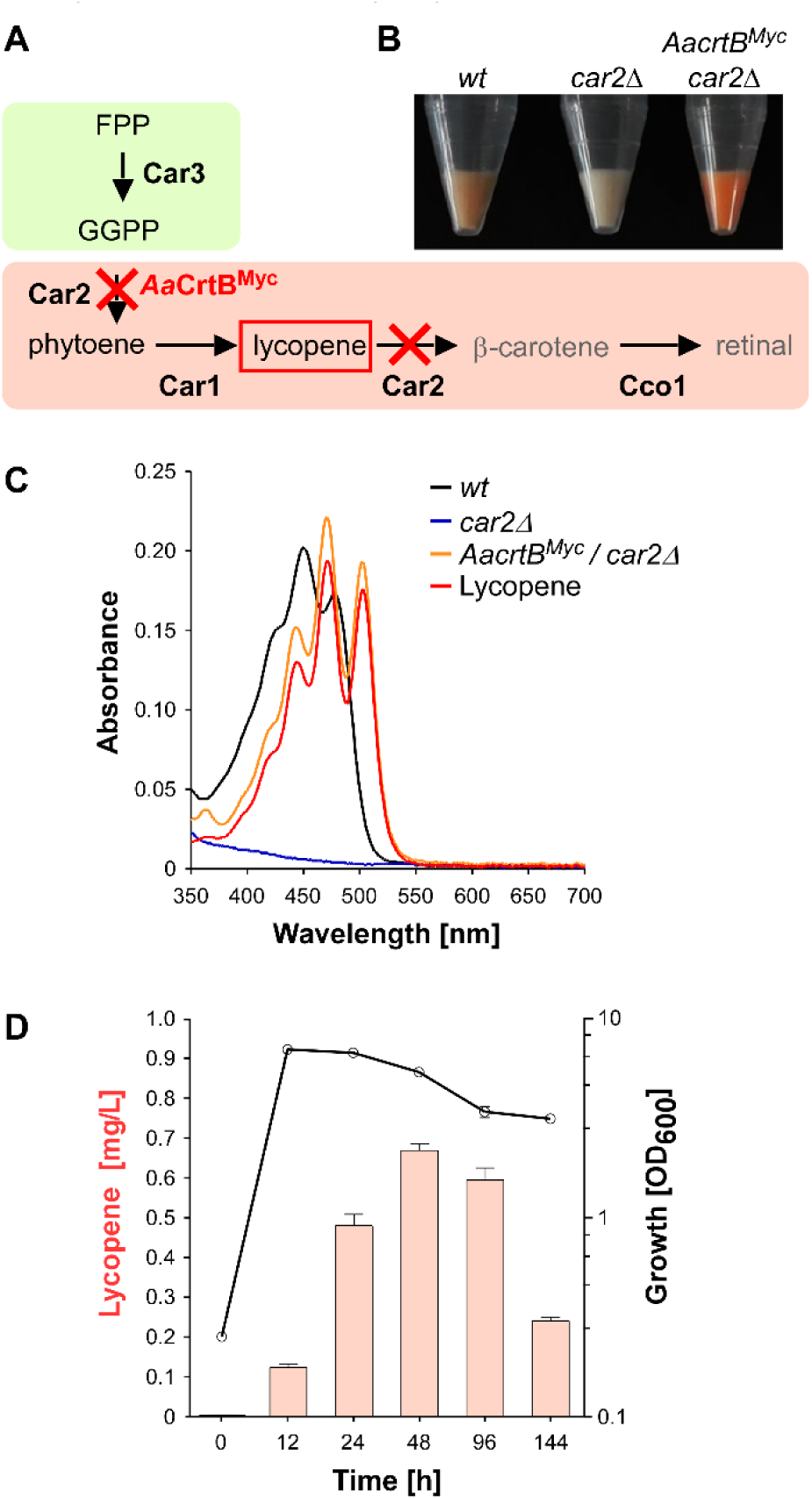
Lycopene production in U. maydis. **(A)** Schematic representation of the carotenoid module given in Figure 1A (red cross indicates gene deletion; AaCrtB^myc^ is the phytoene synthase from *Agrobacterium aurantiacum* containing a triple Myc epitope tag). **(B)** Cell pellets of strains indicated above the image. **(C)** Absorption spectrum of various *U. maydis* strains (genotype as indicated). **(D)** Analysis of lycopene concentrations (left, orange bars) in relation to the growth phase of strain expressing AaCrtB^Myc^ and carrying a deletion of *car2* (OD_600_, black line). Three independent biological experiments (n=3) were carried out. Error bars indicate standard deviation of the mean (SD).

As mentioned above, the carotenoid module was previously studied in *U. maydis* (Figure 1, 2A). Lycopene is naturally produced by desaturation of phytoene, and then further converted in two steps, i.e. cyclisation forming β-carotene and the cleavage of this intermediate to retinal. Car1 is the phyotoene desaturase (Estrada et al., 2009) and deletion of *car1* resulted in loss of carotenoid accumulation (Supplementary Figure S2A-B). Notably, in fungi, Car2 is a bifunctional enzyme, serving as phytoene synthase and lycopene cyclase. In preparation of lycopene production, we deleted *car2* in the wild type, which also abolished carotenoid accumulation due to the bifunctionality of the encoded enzyme (Figure 2A-C, Supplementary Figure S2A-B). The resulting strains exhibited no growth defects. This is consistent with the observation that *cco1* strains and opsins are dispensable for normal growth of *U. maydis* (Estrada et al., 2009). In order to synthesise lycopene in the *car2Δ* strain, we expressed a heterologous phytoene synthase from *Agrobacterium aurantiacum* (*Aa*CrtB; Chen et al., 2016). This should enable heterologous reconstruction of the process and efficient visual detection of the red colour of lycopene (Figure 2A-B). The corresponding bacterial open reading frame was codon-optimised for *U. maydis* (see Materials and Methods) and expression was controlled by the constitutively active promoter P*rpl40*. The respective promoter region was derived from *rpl40*, encoding a ribosomal protein of the large subunit.

The resulting construct was inserted at the defined *upp3* locus of the *car2Δ* strain by homologous recombination. In order to confirm expression of the full length protein, a triple Myc epitope tag was fused at the C-terminus. The expression was verified by Western blot analysis (Supplementary Figure S3A, S4A).

Analysing the resulting strain demonstrated the production of lycopene (Figure 2B-C). Recording an absorption spectrum of cell extracts in *n*-hexane showed that the spectrum was shifted in comparison to the wild type from β-carotene-to lycopene-specific maxima (λ_max_ 450 nm and 503 nm, respectively; Fish et al., 2002), as measured with a commercially available lycopene standard (see Materials and Methods). Studying the production in shake flasks over time revealed that the lycopene titre increased during cell proliferation. Lycopene was still produced in the stationary phase and maximal amounts were detected after 48 hours of culturing (Figure 2D; 0.7 mg/L; see Materials and methods). Expression of *Aa*CrtB^Myc^ in a *car1* and *car2* double deletion strain resulted in no production of carotenoids and confirmed the necessity of Car1 for implementation of lycopene as end product in our strategy (Supplementary Figure S2B). In summary, heterologous expression of a bacterial phytoene synthase in a genetically engineered strain resulted in the efficient production of lycopene as molecular read-out for intracellular FPP levels.

### Metabolic engineering of the mevalonate pathway monitored by lycopene production

In order to increase the activity of the metabolic pathways leading to higher FPP levels, we altered the expression of three biosynthetic genes that were known to encode enzymes with limiting activity in other well-studied systems (Nielsen and Keasling, 2016;Ye et al., 2016): Aat1, Hmg1 and Idi1 (Figure 1A; Table 1). In the case of Hmg1, we generated an N-terminal truncated version of the reductase designated Hmg1^NΔ1-932^. This enzyme is known to be a rate-limiting enzyme in other organisms, whose expression is under tight control (DeBose-Boyd, 2008). It contains a targeting peptide in the N-terminal extension for insertion into the ER membrane and deletion of this region resulted in cytoplasmic localisation and higher activity (Donald et al., 1997;Polakowski et al., 1998;Kampranis and Makris, 2012; truncation indicated in Supplementary Figure S1). In contrast to *S. cerevisiae*, *U. maydis* contains a single Hmg1 enzyme (Table 1) and its activity was already investigated *in E. coli* (Croxen et al., 1994).

To generate *U. maydis* strains with different levels of gene expression, we chose the *ip*^*s*^ locus (Loubradou et al., 2001). Expression was controlled by the strong constitutively active promoter P_*otef*_ and the respective open reading frames were fused at their N-terminus with a triple HA epitope tag for detection (Brachmann et al., 2004). Transformation of linearised plasmids results in two types of homologous recombination events: (i) single or (ii) multiple insertions (Figure 3A). The type of homologous insertion was verified by Southern blot analysis (Figure 3B). In order to verify protein amounts, we performed Western blot analysis and, as expected, multiple insertions resulted in higher protein amounts (Figure 3C). In the case of Hmg1^NΔ1-932HA^, we obtained only single insertion events, although a sufficient number of transformants was screened. This suggests that a strong over-expression of a truncated Hmg1 interferes with growth.

**Figure 3.**
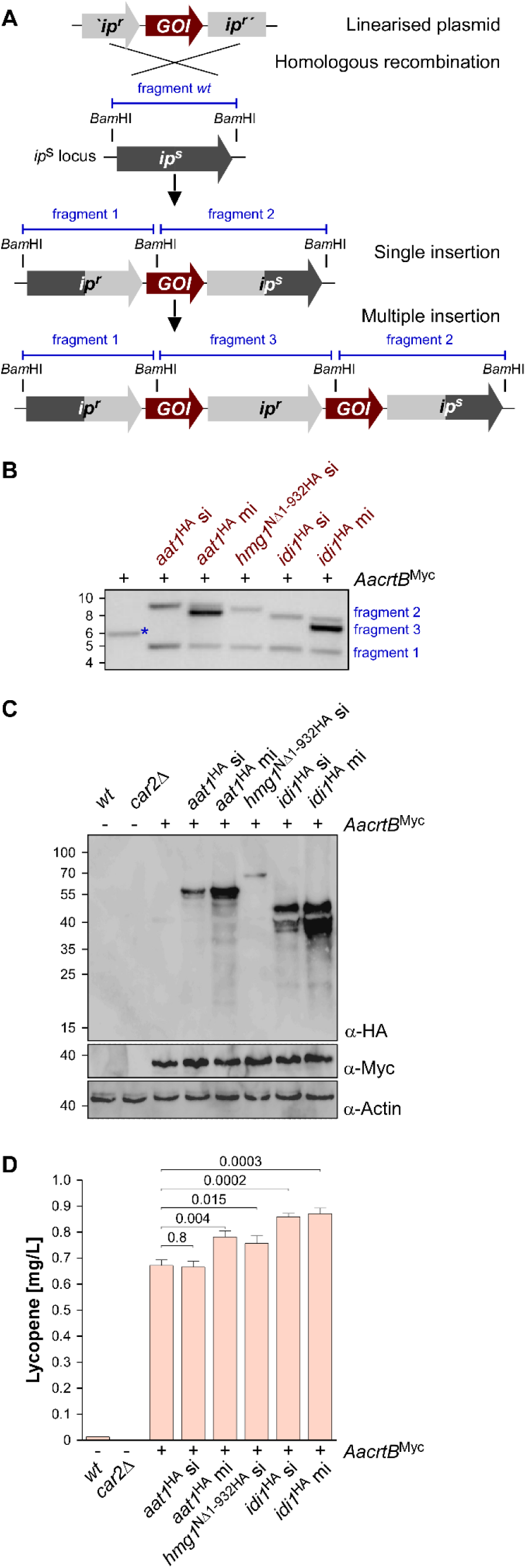
Genetic engineering of the mevalonate pathway. **(A)** Graphical representation of the genetic modification at the *ip*^*s*^ locus encoding an iron sulfur protein conferring carboxin resistance. The wild type (*wt*) version encoding a sensitive version *ip*^*s*^ is given in dark grey. The corresponding resistant version is given in light grey (*ip*^*r*^). The gene of interest (GOI) and fragments expected in Southern blot analysis (shown in B) are given in red and blue, respectively. **(B)** Southern blot analysis of strains indicated above the lanes. The wild type band is indicated by a blue asterisk (size marker in kB, left). **(C)** Western blot analysis of strains indicated above the lanes. The expected molecular weight is 47 kDa for Aat1^HA^, 58 kDa for Hmg1^NΔ1-932HA^, 34 kDa for Idi1^HA^ 37 kDa for AaCrtB^Myc^ and 42 kDa for actin (UMAG_11232). Antibodies are given in the lower right corner (size marker in kDa, left). Note, that the AaCrtBy^Myc^ expressing strains carried a deletion of *car2*. **(D)** Analysis of lycopene concentrations in strains given at the bottom. Three independent biological experiments (n=3) were carried out. Error bars indicate standard deviation of the mean (SD). Statistical significance was calculated using the unpaired two-tailed *t* test and *p*-values were indicated above. Note, that the AaCrtB^Myc^ expressing strains carried a deletion in *car2*.

Assaying the lycopene concentration after 48 hours of incubation in shake flasks revealed in all cases of additional expression of Aat1^Ha^, Hmg1^NΔ1-932HA^ and Idi1^HA^, a statistically significant increase (Figure 3D). In the case of Aat1^HA^ expressing strains, multiple insertions led to a higher lycopene production than a single insertion, indicating that mRNA and protein amounts were limiting. In the case of Hmg1^NΔ1-932HA^, we observed a slight increase in lycopene yield (Figure 3D). A titre of up to 0.9 mg/L could be achieved in strains expressing Idi1^HA^. The amount of lycopene in the strain with multiple insertions of *idi1*^HA^ was not higher, indicating that most likely the enzyme activity, not the protein amount, is limiting or that this was not the limiting step in the strain (Figure 3D, see Discussion). Thus, by addressing known bottlenecks of the mevalonate pathway, we were able to alter terpenoid production, which was easily measured as lycopene production. Hence, lycopene is a good indicator and an efficient as well as robust read-out system for tuning the precursor biosynthetic pathway.

### The subcellular localisation of enzymes involved in FPP synthesis

In other organisms, it has been reported that the mevalonate pathway is compartmentalised. In plants, for example, Aat1 homologues are found in peroxisomes as well as the cytoplasm (Wang et al., 2017). To study the subcellular localisation of the respective enzymes in *U. maydis*, we fused Gfp at their N-termini (enhanced version of the green fluorescent protein, Clontech). We inserted the respective genes at the *ip*^s^ locus and used the constitutively active promoter P_*tef*_ for expression (see Supplementary Table S1). Measuring green fluorescence revealed that the expression of Gfp fusion proteins was detectable (Figure 4A). In Western blot analysis, we verified the expected protein size of the full-length fusion proteins (Supplementary Figure S2C).

**Figure 4.**
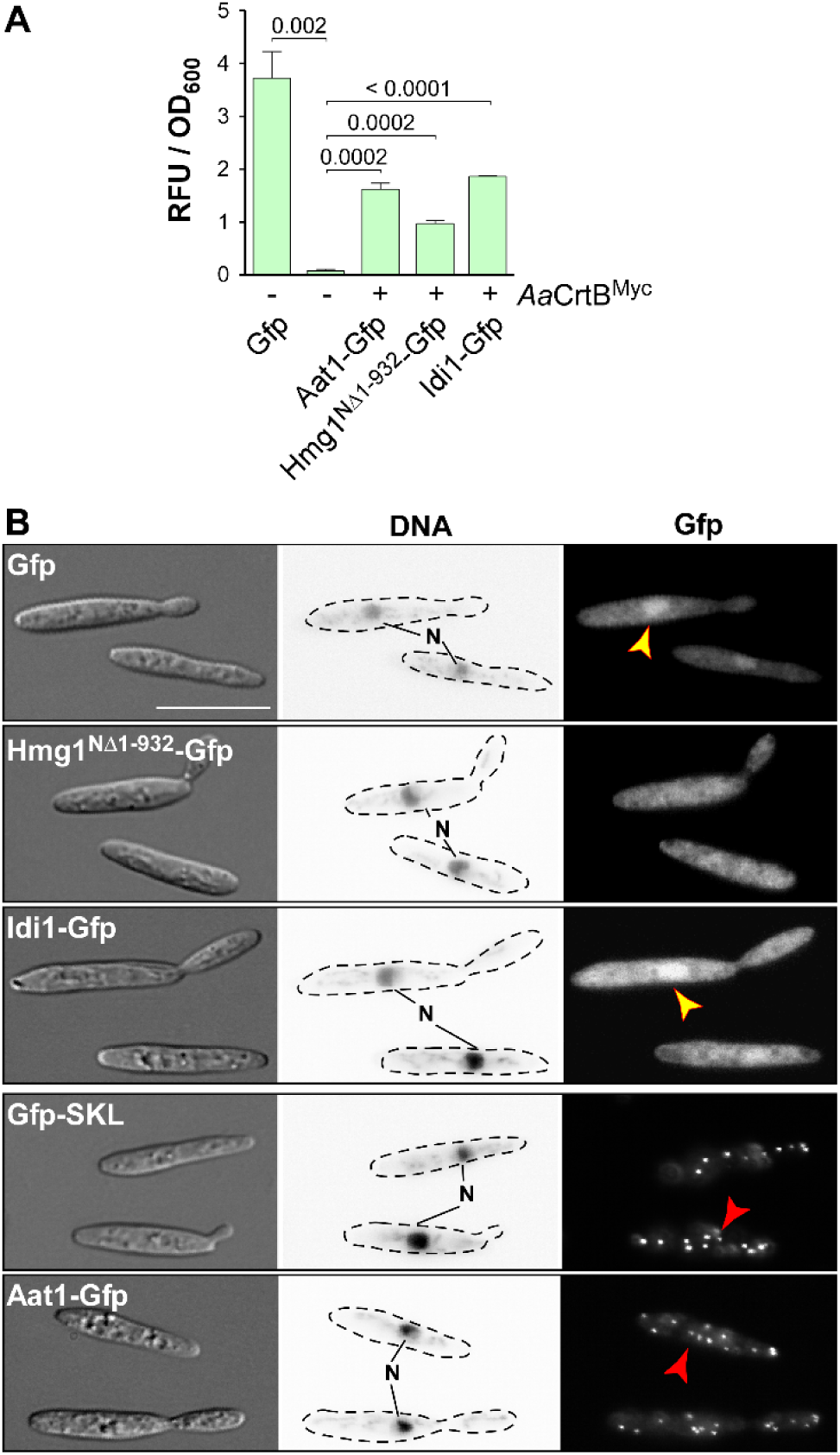
Subcellular localisation of enzymes of the mevalonate pathway. **(A)** Quantification of Gfp expression using fluorimeter measurements. Relative fluorescence units are given relative to the optical density (OD_600_). At least three independent biological experiments (n=3) were performed with three technical replicates per strain. Error bars indicate standard error of the mean (SEM). Statistical significance was calculated using the unpaired two-tailed *t* test and *p*-values were indicated above. Note, that the AaCrtB^Myc^ expressing strains carried a deletion of *car2*. **(B)** Microscopic analysis showing DIC images of fixed cells on the left (size bar, 10 μm). Corresponding staining of DNA with Hoechst 33342 (middle panel; N, nucleus; inverted image) and green fluorescence (Gfp) on the right (yellow and red arrowheads indicate nuclei and peroxisomes, respectively).

Fluorescence microscopy showed that the truncated version Hmg1^NΔ1-932^-Gfp, missing the predicted ER membrane spanning region, localised mainly in the cytoplasm, like unfused Gfp. Idi1-Gfp also exhibited mainly cytoplasmic localisation (Figure 4B). Thus, in contrast to plants (Simkin et al., 2011), geranyl pyrophosphate (GPP) appears to be synthesised in the cytoplasm. Idi1-Gfp also localised to a certain extent to the nucleus (Figure 4B). The same holds true for Gfp and it is known that a small proportion of Gfp and small Gfp fusion proteins mislocalise to the nucleus in *U. maydis* (Figure 4B, see below).

Microscopic observation of Aat1-Gfp revealed the accumulation of fluorescence signals in distinct foci (Figure 4C). This localisation pattern is specific for peroxisomes, as indicated by the localisation of Gfp carrying a C-terminal SKL localisation signal or of peroxisomal protein Pex3-Gfp (Figure 4C, Supplementary Figure S2D).

A second characteristic for peroxisomes in *U. maydis* is their microtubule-dependent transport during hyphal growth (Supplementary Figure S2E; Guimaraes et al., 2015). Fluorescence signals of Aat1-Gfp moved processesively along the hypha as is known for peroxisomes (comparison with movement of Gfp-SKL shown in Supplementary Figure S2E). Thus, either the initial reaction of the mevalonate pathway takes place in peroxisomes or an alternative enzyme is acting during FPP synthesis in the cytoplasm (see Discussion). In essence, studying the subcellular localisation of enzymes of the mevalonate module is highly informative to devise future strategies for pathway engineering.

### Recombinant production of the plant sesquiterpenoid (+)-valencene in *U. maydis*

As proof of principle for sesquiterpenoid production in *U. maydis*, we chose to produce (+)-valencene via heterologous expression of the plant (+)-valencene synthase CnVS, converting FPP to (+)-valencene (Figure 1A; Troost et al., 2019). For efficient detection of protein expression, we fused CnVS with Gfp at the N-terminus, as C-terminal fusions were reported to affect enzyme activity (Kampranis and Makris, 2012). We used the constitutively active promoter P_*otef*_ and inserted the gene at the *upp3* locus of a strain expressing *Aa*CrtB^Myc^ and carrying a deletion in *car2* (see Supplementary Table S1). Expression was verified by fluorimeter measurements and Western blot analysis (Figure 5A, Supplementary Figure S3A).

**Figure 5.**
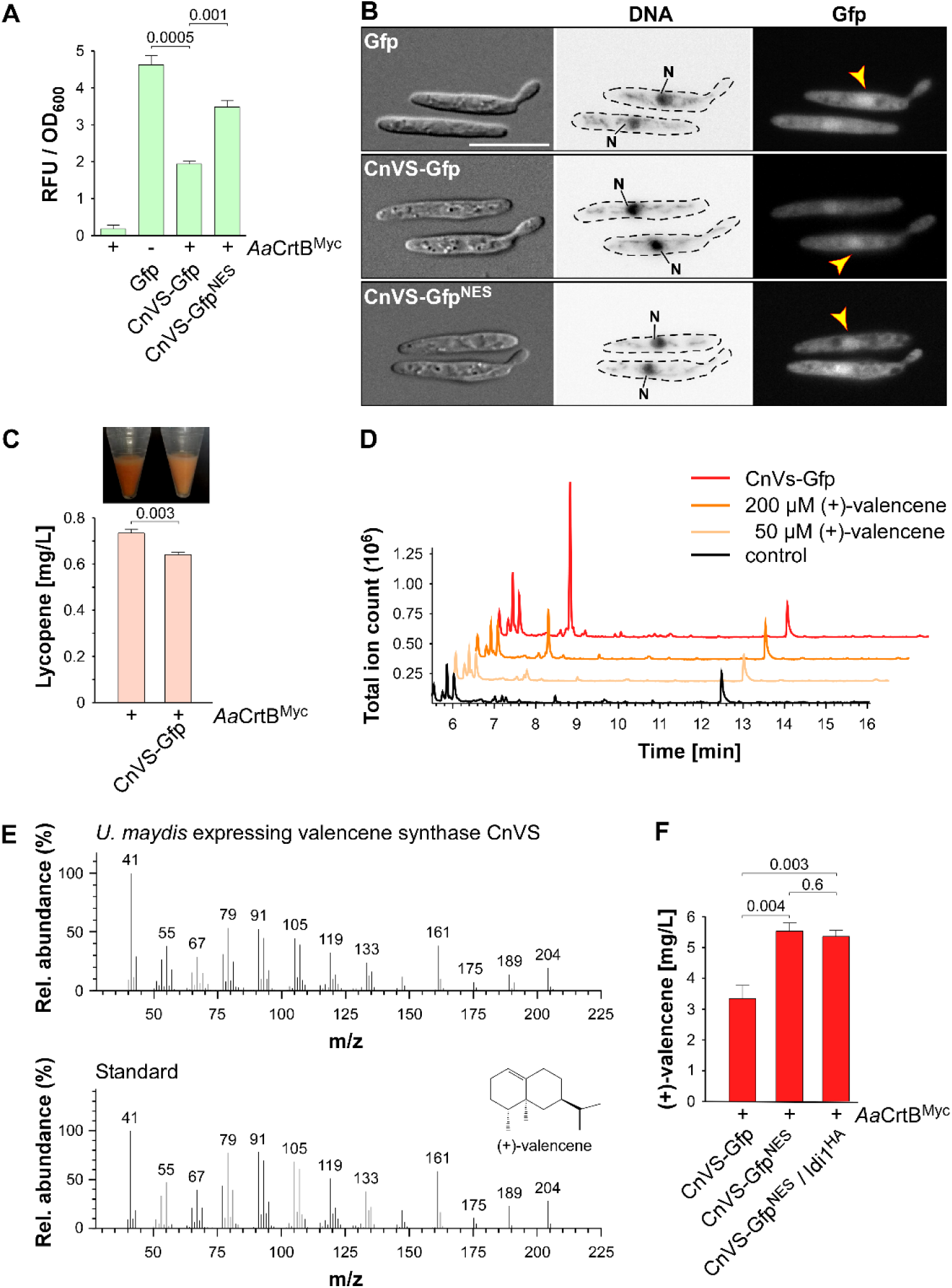
(+)-valencene synthesis in *U. maydis*. **(A)** Quantification of Gfp expression using fluorimeter measurements. Relative fluorescence units are given relative to the optical density (OD_600_). At least three independent biological experiments (n=3) were performed with three technical replicates per strain. Error bars indicate standard error of the mean (SEM). Statistical significance was calculated using the unpaired two-tailed *t* test and *p*-values were indicated above. Note, that the AaCrtB^Myc^ expressing strains carried a deletion of *car2*. **(B)** Microscopic analysis showing DIC images of fixed cells on the left (size bar, 10μm). Corresponding staining of nuclear DNA with Hoechst 33342 (middle panel; N, nucleus; inverted image) and green fluorescence (Gfp) on the right (yellow arrowheads indicate nuclei). **(C)** Cell pellets and lycopene concentrations (orange bars) of strains given at the bottom. Three independent biological experiments (n=3) were carried out. Error bars indicate standard deviation of the mean (SD). Statistical significance was calculated using the unpaired two-tailed *t* test and *p*-values were indicated above. Note, that the AaCrtB^Myc^ expressing strains carried a deletion of *car2*. **(D)** GC-MS chromatogram of (+)-valencene from CnVS expressing strain and the corresponding standard diluted in n-dodecane samples of the negative control. **(E)** Fragmentation pattern of peaks at 7.3 min (shown in D). **(F)** Concentration of (+)-valencene produced in the culture, determined by GC-FID according to commercial reference compound (Supplementary Figure S3B). Three independent biological experiments (n=3) were carried out. Error bars indicate standard deviation of the mean (SD). Statistical significance was calculated using the unpaired two-tailed *t* test and *p*-values were indicated above. Note, that the AaCrtB^Myc^ expressing strains carried a deletion of *car2*.

Studying the subcellular localisation showed the presence of CnVS-Gfp in the cytoplasm in order to ensure substrate access, but also to some extend in the nucleus (Figure 5B). To enhance cytoplasmic localisation, we fused a nuclear export signal (NES) from murine minute virus (MTKKFGTLTI; Engelsma et al., 2008) to the N-terminus of Gfp resulting in CnVS-Gfp^NES^. Fluorescence microscopy revealed that the protein was also expressed in this form but its nuclear localisation was only slightly reduced (Figure 5B). Hence, the heterologous NES did not function efficiently in *U. maydis*. However, we observed that the protein amount of CnVS-Gfp^NES^ was significantly higher than CnVS-Gfp (Figure 5A; Supplementary Figure S3A). The N-terminal NES sequence most likely improved expression or protein stability as a side effect.

Measuring the lycopene titre of the strain co-expressing CnVS-Gfp and *Aa*CrtB^Myc^ (*car2* background) showed a significant decrease of lycopene compared to the progenitor strain, which can also be detected by visual inspection (Figure 5C). Apparently, a proportion of the FPP is no longer available for lycopene synthesis, suggesting the functionality of the heterologous CnVS-Gfp. In order to measure the volatile (+)-valencene, we incubated strains with *n*-dodecane to collect the product (see Material and Methods). After 48 hours of incubation in shake flasks, we measured (+)-valencene in the *n*-dodecane phase by gas chromatography (Figure 5D). We observed a prominent peak in GC-MS analysis with identical retention time of 7.3 min to the commercial reference (+)-valencene. Importantly, this peak was absent in the negative control strain (Figure 5D). Furthermore, analysing a fragmentation pattern in mass spectrometry (MS) identified a pattern of peaks characteristic for (+)-valencene (Troost et al., 2019). Thus, the identified substance is most likely the desired product. Using a commercial reference (Merck), we generated a standard calibration curve to quantify the titre via GC-FID analysis (Supplementary Figure S3B). Up to 5.5 mg/L (+)-valencene could be obtained from the CnVS-GfpNES expressing strain (*Aa*CrtB^Myc^/*car2Δ*; Figure 5F).

In order to verify whether the N-terminal fusion of Gfp interferes drastically with the enzyme activity, we compared the (+)-valencene titre from a CnVS-Gfp expressing strain (3.3 mg/L) with that of a strain expressing an untagged version (4.8 mg/L; Supplementary Figure S3C). Hence, the Gfp fusion only slightly reduced the enzyme activity. However, it would be advisable to use an untagged CnVS for an improved production strain.

Finally, we tested whether overexpression of Idi1^HA^, which improved the carbon flux within the mevalonate pathway (see above), can improve the titre of (+)-valencene. To this end, the corresponding gene was inserted at the *cco1* locus of the strain expressing CnVS-Gfp^NES^. For *idi1*^*HA*^ expression, we used the constitutively active promoter P_*rpl10*_. The promoter region was derived from *rpl10*, encoding ribosomal protein 10 of the large subunit.

The lycopene titre was increased to 0.9 mg/L (Supplementary Figure 3E), which is comparable to the values obtained when Idi1^HA^ was expressed at the *ip*^*s*^ locus (Figure 3D). However, (+)-valencene production was not increased in this strain (Figure 5F). For future attempts, it might be advantageous to downregulate expression of *Aa*CrtB^Myc^ or Car3 to redirect more FPP to sesquiterpenoids (see Discussion). In essence, we succeeded in the production of the widely produced plant sesquiterpenoid (+)-valencene, which was chosen here as a common model compound, by reengineering the intrinsic FPP pathway.

### Recombinant production of the basidiomycete sesquiterpenoid α-cuprenene in *U. maydis*

As pointed out above, the basidiomycete *U. maydis* might serve as a suitable production platform for the synthesis of sesquiterpenoids from higher basidiomycetes. For this purpose, we chose to express the sesquiterpenoid synthase Cop6 from the mushroom *C. cinerea* in *U. maydis* (Agger et al., 2009). We fused Gfp or Gfp^NES^ to the N-terminus of Cop6, used the constitutively active promoter P_*otef*_ and targeted the corresponding construct to the *upp3* locus of a strain expressing *Aa*CrtB^Myc^ and carrying a deletion of *car2* (see Supplementary Table S1). The expression was confirmed via fluorimeter measurements and Western blot analysis (Figure 6A; Supplementary Figure S4A). Analysis of the subcellular localisation revealed that Cop6-Gfp localised to the cytoplasm (Figure 6B). Consistent with the aforementioned localisation of CnVS-Gfp^NES^, the NES version did not prevent nuclear accumulation but resulted in higher protein amounts (Figure 6A-B, Supplementary Figure S4A).

**Figure 6.**
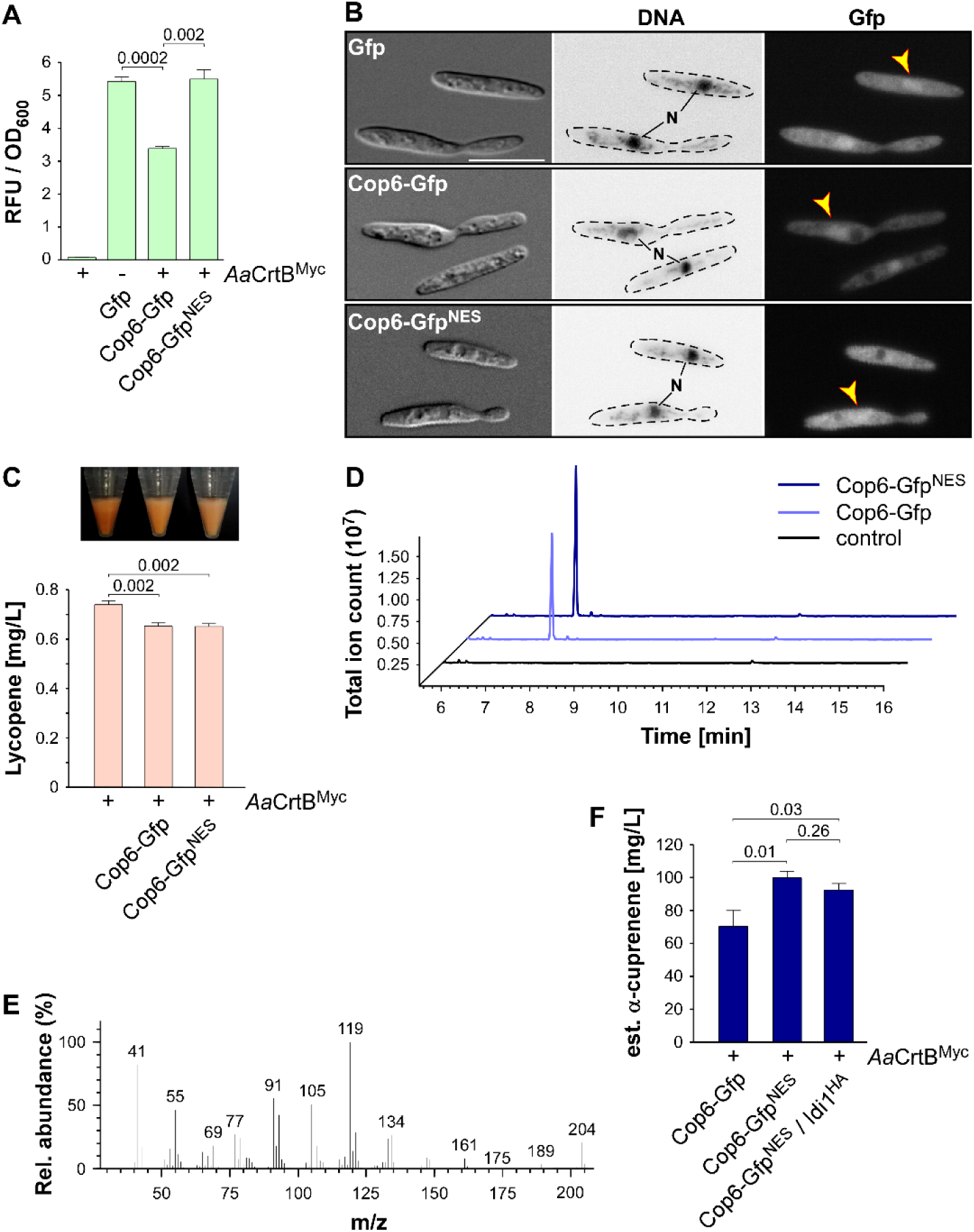
α-cuprenene synthesis in *U. maydis*. **(A)** Quantification of Gfp expression using fluorimeter measurements. Relative fluorescence units (RFU) are given relative to the optical density (OD_600_). At least three independent biological experiments (n=3) were performed with three technical replicates per strain. Error bars indicate standard error of the mean (SEM). Statistical significance was calculated using the unpaired two-tailed *t* test and *p*-values were indicated above. Note, that the AaCrtB^Myc^ expressing strains carried a deletion of *car2*. **(B)** Microscopic analysis showing DIC images of fixed cells on the left (size bar, 10 μm). Corresponding staining of DNA Hoechst 33342 (middle panel; N, nucleus, inverted image) and green fluorescence (Gfp) on the right (yellow arrowheads indicate nuclei). **(C)** Cell pellets and lycopene concentrations (orange bars) of strains given at the bottom. Three independent biological experiments (n=3) were carried out. Error bars indicate standard deviation of the mean (SD). Statistical significance was calculated using the unpaired two-tailed t test and p-values were indicated above. Note, that the AaCrtB^Myc^ expressing strains carried a deletion of *car2*. **(D)** GC-MS chromatogram of α-cuprenene from Cop6 expressing strains. **(E)** Fragmentation pattern of peaks at 7.3 min given in (D). **(F)** Estimated concentration of α-cuprenene determined via GC-FID using the reference compound β-caryophyllene (Supplementary Figure S4B). Three independent biological experiments (n=3) were carried out. Error bars indicate standard deviation of the mean (SD). Statistical significance was calculated using the unpaired two-tailed *t* test and *p*-values were indicated above. Note, that the AaCrtB^Myc^ expressing strains carried a deletion of *car2*.

As was the case with expression of CnVS described above, we observed that the amount of lycopene was reduced in strains co-expressing the sesquiterpenoid synthase and *Aa*CrtB^Myc^ (*car2Δ* background), suggesting that part of the FPP was redirected to the recombinant sesquiterpenoid (Figure 6C). Respective strains were incubated for 48 hours in shake flasks and the α-cuprenene was trapped in *n*-dodecane (see Materials and methods). The GC-MS analysis of the organic phase showed an additional peak at a retention time of 7.4 min that was absent in the negative control (Figure 6D). In order to confirm α-cuprenene production, we analysed the fragmentation pattern of this peak in mass spectrometry (Figure 6E). The fragmentation pattern was consistent with reported data for α-cuprenene (Agger et al., 2009). The total ion count was slightly higher in the NES version, most likely due to the higher enzyme amount in the Cop6-Gfp^NES^ expressing strain (Figure 6A-B). Finally, we estimated the yield of α-cuprenene production. Due to the absence of a commercial reference for α-cuprenene, we generated a standard calibration curve with the similar reference sesquiterpenoid β-caryophyllene (Merck) and compared the intensity of the electrically charged particles via GC-FID (Supplementary Figure S4B). We observed the highest amount of α-cuprenene in the Cop6-GfpNES expressing strain and estimated a titre of 0.1 g/L (Figure 6F). Overexpression of Idi1^HA^ at the *cco1* locus did not increase the titre further (Figure 6F; see Discussion). In summary, we also succeeded in synthesising the fungal sesquiterpenoid α-cuprenene, in addition to the plant-derived (+)-valencene, demonstrating that *U. maydis* serves as a promising novel host for the production of such specific sesquiterpenoids.

## Discussion

In this study, we present a straightforward strategy to engineer the terpenoid metabolism and produce the carotenoid lycopene as well as plant and fungal sesquiterpenoids in *U. maydis*. We have established lycopene production as an efficient read-out for the activity of the FPP-dependent pathway. This allowed initial pathway engineering and the production of plant (+)-valencene and fungal α-cuprenene as proof of principle.

### Establishing lycopene production as molecular read-out, reflecting internal FPP levels

To establish a terpenoid producing strain, we addressed early on the establishment of a simple read-out system for the activity of the FPP-based metabolic network. To this end, we redirected the intrinsic carotenoid pathway towards the production of lycopene by deletion of *car2* and heterologous expression of the phytoene synthase *Aa*CrtB (Chen et al., 2016). Importantly, loss of retinal biosynthesis does not affect the growth of cells and currently, no light-regulated biological function could be assigned to retinal-dependent opsins (Estrada et al., 2009). Our strategy enabled visual and quantitative detection of the carotenoid lycopene, which was used for initial pathway engineering (see below). Genetic engineering that enhanced FPP synthesis, like overexpression of Idi1, resulted in increased lycopene levels. Conversely, expression of sesquiterpenoid synthases consuming FPP reduced lycopene amounts. Thus, the intracellular FPP levels correlated with lycopene yields.

Similar strategies have been followed in *S. cerevisiae*, where GGPP was used as a metabolic branching point to synthesise carotenoids. The resulting strains were successfully applied to improve terpenoid production by carotenoid-based screening of mutants or by automated lab evolution (Ozaydin et al., 2013;Reyes et al., 2014;Trikka et al., 2015). Redirecting naturally occurring carotenoid pathways towards sesquiterpenoids was successfully achieved in *Corynebacterium glutamicum* and the red yeast *Xanthophyllomyces dendrorhous*. Both microorganisms are, in contrast to *S. cerevisiae*, natural producers of carotenoids (Melillo et al., 2013;Frohwitter et al., 2014).

In order to advance the system in the future, controlled down-regulation of GGPP synthase Car3 expression will reduce its activity and channel higher amounts of FPP to terpenoid production. Regulated expression is advantageous since high FPP concentrations are toxic for microorganisms and increased pathway activity resulting in higher FPP levels could even limit production (Dahl et al., 2013). Thus, lycopene production serves as a safety valve for excess FPP. In *E. coli* this problem was addressed by using stress responsive promoters that respond to high FPP levels. The identified promoters were used to create a functional FPP biosensor and improved the production of amorphadiene (Dahl et al., 2013). Finally, besides the use of lycopene as a read-out system, *U. maydis* might offer the possibility to serve as an alternative host for lycopene production. Such a sustainable biotechnological approach prevents the use of nutrient-rich food like tomatos for lycopene extraction and avoids the risk of contamination by bacterial toxins when produced, for example, in *E. coli*. The lycopene production titre of our current system is rather low. However, pathway engineering in other fungal microorganisms like *S. cerevisiae* resulted in strains producing 0.3 g/L of lycopene in shake flask fermentation (Shi et al., 2019). Thus, comparable pathway engineering in *U. maydis* could result in similar increases of the yield.

### Genetic engineering for higher biosynthesis of FPP

Initially, we carried out a bioinformatics analysis allowing us to annotate the mevalonate, prenyl phosphate and carotenoid modules. As pointed out above we started metabolic engineering of the mevalonate pathway at three different points: overexpression of Aat1, Idi1 and a truncated version of Hmg1, containing only the catalytic domain (Kampranis and Makris, 2012).

In each case, the accumulation of lycopene was higher, supporting the accuracy of our pathway prediction. The most successful approach was transcriptional upregulation of *idi1*, indicating that this key step is also a clear bottleneck of the pathway in *U. maydis*. Idi1 is also known as the bottleneck in other systems like *S. cerevisiae*, where it was also initially tackled by *IDI1* overexpression or expression of heterologous plant enzymes with better performances (Ignea et al., 2011;Ye et al., 2016). Multiple insertions of the Idi1 expression construct in *U. maydis*, however, did not increase lycopene levels further, suggesting that the Idi1 mRNA amount is no longer the limiting factor.

Expression of the truncated version of Hmg1 was inspired by work in *S. cerevisiae*, where removing the N-terminal domain resulted in cytoplasmic localisation due to detachment of the ER membrane and increased protein stability (Donald et al., 1997;Kampranis and Makris, 2012). Consistently, overexpression of Hmg1^N1Δ-932^ resulted in higher lycopene amounts. In this case, we did not obtain multiple insertions of the *hmg1*^NΔ1-932^ allele at the *ips* locus, suggesting that high levels of Hmg1^NΔ1-932^ cause growth defects. In the future, these alterations should be combined in a single production strain to further improve the performance of the underlying metabolic network.

Importantly, we verified the subcellular localisation of the enzymes. Idi1 and the truncated Hmg1 version both localised in the cytoplasm as expected. But to our surprise Aat1 exhibited a peroxisomal localisation, although no conventional peroxisomal targeting sequences were detectable. Peroxisomal targeting of acetoacetyl-CoA thiolases is reminiscent of the subcellular localisation in mammalian and plant cells (Kovacs et al., 2007;Simkin et al., 2011). It was proposed in mammalian cells that the complete mevalonate pathway is occurring in peroxisomes (Kovacs et al., 2007). Since we observed that Idi1 appears to be cytoplasmic in *U. maydis*, the late steps of the pathway most likely take place in the cytoplasm. Thus, in basidiomycetes, the subcellular compartmentalisation of the mevalonate pathway might be different. Interestingly, there are four additional enzymes annotated as putative acetoacetyl-CoA thiolases in the proteome of *U. maydis* (UMAG_03298, 01843, 01090 and 02715). In principle, these enzymes could be participating in the mevalonate pathway in the cytoplasm. In essence, successfully carrying out the initial steps in pathway engineering, we were able to lay a solid foundation for future improvements.

### Production of plant and fungal sesquiterpenoids in *U. maydis*

In order to demonstrate sesquiterpenoid production, we chose plant (+)-valencene and fungal α-cuprenene to be synthesised in *U. maydis*. (+)-valencene is extensively used in the flavour and fragrance industries, but also shows antagonistic activity against the plant pathogenic nematode *Heterodera schachtii* (Troost et al., 2019; Schleker et al. manuscript in preparation). High titres of 352 mg/L and 540 mg/L were reached in optimised *Rhodobacter sphaeroides* and *S. cerevisiae* strains, respectively (Beekwilder et al., 2014;Chen et al., 2019). Currently, a sustainable production process is marketed by the companies Evolva and Isobionics (now BASF) using the aforementioned microorganisms (Schempp et al., 2018;Chen et al., 2019). We therefore chose this target compound as a very prominent representative of heterologously produced sesquiterpenoids to benchmark our approach. Expression of the plant (+)-valencene synthase CnVS in *U. maydis* alone resulted in (+)-valencene levels of 5.5 mg/L in shake flask fermentation, without extensive pathway engineering. To put it into perspective, heterologous expression of *CnVS* in *S. cerevisiae* resulted in initial titres of 1.3 mg/L (Cankar et al., 2014). Thus, our initial yield of (+)-valencene in *U. maydis* is a promising starting point but definitely needs further improvement.

α-cuprenene is originally synthesised by the sesquiterpenoid synthase Cop6 from *Coprinopsis cinerea*. The corresponding gene is part of a mini gene cluster flanked by *cox1* and *cox2* encoding cytochrome P450 monooxygenases. α-cuprenene is converted by these enzymes to lagopodin B that has antibacterial activity against Gram-positive bacteria (Agger et al., 2009;Stöckli et al., 2019). Interestingly, all members of the cluster are transcriptionally activated after co-cultivation with Gram-positive bacteria (Stöckli et al., 2019). We chose this target compound as a representative of basidiomycete specialty chemicals that are not yet widely accessible to test applicability of our approach. We succeeded in the production of α-cuprenene in *U. maydis* in reasonable amounts, reaching the estimated titre of 0.1 g/L in shake flask fermentation. Consistently, α-cuprenene was also produced in the basidiomycete yeast *X. dendrorhous*, using a similar strategy where Cop6 was expressed to redirect FPP towards α-cuprenene. A comparable titre of 0.08 g/L was obtained after four days of cultivation and strains were able to produce α-cuprenene and astaxanthin simultaneously, indicating that, like in our case, the wild type FPP pool was not limited in sesquiterpenoid synthesis (Melillo et al., 2013).

In the future, Cop6 may be co-expressed with the cytochrome P450 monooxygenases Cox1 and Cox2 for the production of lagopodin B in *U. maydis* (Stöckli et al., 2019). Notably, the titre of the fungal compound was ten times higher than the plant compound. This is consistent with our initial hypothesis that the basidiomycete *U. maydis* might serve as a promising expression host for the production of sesquiterpenoids from higher basidiomycetes.

### Conclusion and outlook

We followed the microbial cell factory concept to establish the model basidiomycete fungus *U. maydis* as a novel chassis for the production of a broad range of terpenoids. As proof of concept, we have successfully produced two different sesquiterpenoids in *U. maydis* by modification of the existing mevalonate pathway and introduction of heterologous enzymes. In the future, additional sesquiterpenoid synthases from higher basidiomycetes like *Omphalotus olearius* could be expressed to produce illudins and other compounds (Schmidt-Dannert, 2015;Xiao and Zhong, 2016). To increase the yield of the desired compounds, there are several straightforward strategies possible that were successful in other microbial systems (Kampranis and Makris, 2012). Upregulation of the desired pathway would be possible by the expression of heterologous enzymes with higher activity like optimised versions of bacterial Aat1 (Shiba et al., 2007). Competing pathways, such as that for ergosterol biosynthesis, could be reduced by downregulation of Erg9 squalene synthase expression or by treatment with chemical inhibitors, which block the synthesis of ergosterol (Asadollahi et al., 2010). Alternatively, deletion of *fer4*, encoding an enoyl-CoA hydratase, will prevent the consumption of hydroxymethyl glutaryl-CoA during ferrichrome A synthesis (Winterberg et al., 2010). Finally, the fusion of FPP synthase (FPPS) and germacrene A synthase (GAS) has been shown to increase sesquiterpenoid yield, as FPP was directly funnelled into product formation (Hu et al., 2017). Furthermore, in combination with the enhanced biomass degrading ability of *U. maydis* for use of alternative carbon sources (Geiser et al., 2016b;Stoffels et al., 2020), we envision to generate a sustainable consolidated strategy for next generation bioengineering of terpenoid production.

## Materials and methods

### Bioinformatics analysis

The KEGG database (Kyoto Encyclopedia of Genes and Genomes, https://www.genome.jp/kegg/) was used to identify candidates of the mevalonate pathway enzymes in *U. maydis* (Figure 1, Table 1). For verification, amino acid sequences were compared to known enzymes from well-studied eukaryotes like *S. cerevisiae*, *H. sapiens* and *A. thaliana* using BLASTP (https://blast.ncbi.nlm.nih.gov/blast/). Clustal Omega and GeneDoc 2.6 were used for multiple amino acid sequence alignments and graphical representation (Supplementary Figure S1; Larkin et al., 2007). Domain structure was analysed with SMART (Simple Modular Architecture Research Tool; analysis performed April 2020; Schultz et al., 1998;Letunic and Bork, 2018).

### Plasmids, strains and growth conditions

Standard molecular cloning procedures were carried out as published previously (Brachmann et al., 2004;Pohlmann et al., 2015). In brief, cloning was performed using *E. coli* K-12 derivate Top10 (Life Technologies, Carlsbad, CA, USA) and standard techniques like Golden Gate as well as Gibson assembly were followed (Gibson et al., 2009;Terfrüchte et al., 2014). All *U. maydis* strains were generated by transformation of cells with linearised plasmids and homologous recombination events were verified by Southern blot analysis (Brachmann et al., 2004). Genomic DNA of wild type strain UM521 (*a1b1*) was used as template for PCR. Oligonucleotides used for molecular cloning are listed in Supplementary Table S4. For efficient expression of heterologous genes the codon usage was optimised using online tools from IDT (Integrated DNA Technologies, Leuven, Belgium) in the case of *Aa*CrtB^Myc^ or tailor-made context-dependent codon usage tools (Zarnack et al., 2006; Zhou et al., 2018; http://dicodon-optimization.appspot.com) in the case of CnVS and Cop6. Cognate nucleotide sequences were chemically synthesised by IDT. Proteins were tagged with Gfp (enhanced green fluorescent protein; Clontech, Mountain View, CA, USA) at their N-terminus unless noted otherwise. For recycling of resistance markers, the FLP-FRT system with different FRT site variants was applied as described (Khrunyk et al., 2010).

The cultivation conditions and antibiotics used for *U. maydis* strains are described in Brachmann et al., 2004. In brief, complete medium was supplemented with 1% glucose (CM-glc) and strains were incubated at 28°C shaking in baffled flasks with 200 rpm. In order to induce the filamentous form of laboratory strain AB33, the medium was exchanged from CM-glc to nitrate minimal medium supplemented with 1% glucose (NM-glc). For monitoring cell growth and lycopene accumulation, a pre-culture was diluted to a starting OD600 of 0.25 in 500 mL volume non-baffled flask containing a total of 10 mL culture and incubated in white light (adapted from Estrada et al., 2009). Detailed information on plasmids and strains is given in Supplementary Table S1-S3. Plasmid sequences are available on request.

### Lycopene extraction and quantification

Lycopene extraction was adopted and modified as published (Ukibe et al., 2008). Cell pellets of a 10 mL culture after 48 hours of incubation were washed in double-distilled water before disruption in acetone together with glass beads by shaking twice at a frequency of 30 Hz for 15 min in a pebble mill (MM400, Retsch GmbH, Haan, Germany). Afterwards, each suspension was heated for 10 min at 65°C to extract lycopene. The acetone was completely dried overnight at room temperature and samples were re-dissolved in 1 mL *n*-hexane. To remove cell debris, centrifugation at 16,000 *g* for 10 min was performed twice. The lycopene extraction process was carried out at low light conditions.

A commercial reference of lycopene was purchased (SMB00706, Merck, Darmstadt, Germany) to compare absorbance wavelengths with lycopene extracts. Absorbance scanning was carried out in a UV-Vis spectrophotometer (Genesys 10S UV-Vis, Thermo Fisher Scientific, Waltham, MA, USA) recording from 350 nm to 700 nm. For quantification of lycopene, absorption at 503 nm was determined and titre was assessed using molecular extinction coefficient 172,000 M^−1^cm^−1^ of lycopene in *n*-hexane (Fish et al., 2002).

### Protein extraction and Western blot analysis

A culture of 20 mL growing to an OD600 of ca. 2 was harvested by centrifugation (16,000 *g* for 10 min at room temperature). Cell pellets were frozen in liquid nitrogen and then disrupted in a pebble mill (MM400, Retsch GmbH) using a frequency of 30 Hz three times for one min with steal beads. Afterwards, urea buffer (8 M urea, 50 mM Tris-HCl, pH 8) supplemented with protease inhibitors (1 tablet of complete protease inhibitor per 20 mL; Roche, Mannheim, Germany, 0.1 M PMSF and 0.5 M benzamidine) was used for resuspension at 4°C. Cell debris was pelleted by centrifugation at 16,000 *g* for 15 min at 4°C. The protein concentration in the supernatant was determined using a Bradford assay (Bio-Rad, Munich, Germany). 10% SDS-PAGE gels were used for separation of proteins, which were transferred to nitrocellulose membranes (Amersham Protran 0.45 NC Western blotting membrane, GE Healthcare Life Sciences, Munich, Germany) using a semi-dry Western blot procedure. Epitope-tagged proteins were detected using different primary antibodies produced in mouse, α-Gfp (monoclonal, Roche, Freiburg, Germany; 1:1,000 dilution), α-HA (monoclonal, Roche; 1:4,000 dilution), α-Myc (monoclonal, Merck; 1:5,000 dilution) and α-actin (monoclonal antibody raised against actin from chicken gizzard, MP Biomedicals, Eschwege, Germany; 1:500 dilution). Secondary α-mouse IgG-HRP conjugate (Promega, Mannheim, Germany; 1:4,000 dilution) was used for detection. The developing step was performed with Amersham ECL prime detection reagent and a LAS4000 chemiluminescence imager (both GE Healthcare Life Sciences).

### Fluorimetric measurements

A pre-culture was diluted to an OD_600_ 0.5 in CM-glc for all strains and 1 mL of each culture was harvested at 16,000 *g* for 5 min at room temperature. The cell pellets were washed twice in double-distilled water. Afterwards, each cell pellet was resuspended in 1 mL of double-distilled water. 200 µL of each sample was transferred into black 96-well plates (Greiner Bio-One, Frickenhausen, Germany) for measurements in an Infinite M200 plate reader (Tecan Group Ltd., Männedorf, Switzerland). As a blank 200 µL of double-distilled water was used. Within the microplate reader, measurements of OD600 and fluorescence intensity were performed. In the case of Gfp, an excitation wavelength of 483 nm and an emission wavelength of 535 nm were used. At least three independent biological experiments were performed with three technical replicates per strain.

### Fluorescence microscopy

To visualise the nucleus, cells were first fixed with 1% formaldehyde for 30 min and washed twice in PBS. Afterwards, the DNA was stained with Hoechst 33342 dye (H1399, Thermo Fisher Scientific). In brief, 10 mg/mL stock solution was diluted to 1:2000 in PBS and the fixed cells were stained for 10 min at room temperature. Excess dye was washed three times with PBS.

Microscopy was carried out as described before (Baumann et al., 2016;Jankowski et al., 2019) using two systems: (i) a wide-field microscope set-up from Visitron Systems (Puchheim, Germany), Axio Imager M1 equipped with a Spot Pursuit CCD camera (Diagnostic Instruments, Sterling Heights, MI, USA) and the objective lens Plan Neofluar (40 ×, NA 1.3; 63 ×, NA 1.25; Carl Zeiss, Jena, Germany). The excitation of fluorescently labelled proteins was carried out using an HXP metal halide lamp (LEj, Jena, Germany) in combination with a filter set for green fluorescent protein (ET470/40BP, ET495LP and ET525/50BP) and Hoechst 33342 dye (HC387/11BP, BS409LP and HC 447/ 60BP; AHF Analysentechnik, Tübingen, Germany). The microscopic system was controlled by MetaMorph software (Molecular Devices, version 7, Sunnyvale, CA, USA). The program was also used for image processing, including the adjustment of brightness and contrast.

(ii) In order to record fluorescence signals localised in subcellular compartments with higher sensitivity, laser-based epifluorescence microscopy was performed on a Zeiss Axio Observer.Z1 equipped with CoolSNAP HQ2 CCD (Photometrics, Tuscon, AZ, USA) and ORCA-Flash4.0 V2 + CMOS (Hamamatsu Photonics Deutschland GmbH, Geldern, Germany) cameras. The microscopy set-up was the same as described above (Jankowski et al., 2019). For excitation, a VS-LMS4 Laser-Merge-System (Visitron Systems) that combines solid-state lasers for excitation of Gfp (488 nm at 50 or 100 mW) was used. Videos were recorded with an exposure time of 150 ms and 150 frames were taken. All videos and images were processed and analysed using Metamorph (Version 7.7.0.0, Molecular Devices). Kymographs were generated using a built-in plugin.

### Analysis of sesquiterpenoids using GC-MS

For the analysis of (+)-valencene and α-cuprenene, 10 mL of cells grown to an OD600 of approximately 6 were harvested after 48 hours. To trap the secreted sesquiterpenoids, 500 μL of *n*-dodecane (D0968, TCI Deutschland GmbH, Eschborn, Germany) were added to the shaking culture. To ensure phase separation, the *n*-dodecane containing layer was centrifuged twice at 16,000 *g* for 10 min. The *n*-dodecane phase-containing sesquiterpenoids was analysed using a GC/MS-QP2010 (Shimadzu, Tokyo, Japan) equipped with a FS-Supreme-5 column (30 m × 0.25 mm × 0.25 μm; Chromatographie Service GmbH, Langerwehe, Germany). The GC-MS conditions were adopted from a previous study (Schulz et al., 2015). Temperatures of the injector and interface were set at 250°C and 285°C, respectively. The carrier gas was helium and its velocity was set to 30 cm sec^−1^. 1 μL of the sample was injected with a split ratio of 10. The column temperature was sequentially changed and maintained at 130°C for 3 min, ramped to 260°C at a rate of 10°C min−1, held at 260°C for 1 min, ramped to 300°C at a rate of 40°C min^−1^ and held at 300°C for 1 min. In the case of (+)-valencene, a purchased reference compound (75056, Merck) was diluted as 50 μM and 200 μM in *n*-dodecane samples of the negative control. The retention time and fragmentation pattern of the mass spectrum obtained with the *U. maydis* sample were compared with the reference. In addition, the fragmentation patterns of (+)-valencene and α-cuprenene were compared with the previously reported data (Agger et al., 2009;Troost et al., 2019).

### Quantification of sesquiterpenoid production with GC-FID

In order to determine product titres of (+)-valencene and α-cuprenene, *n*-dodecane samples were subjected to the Agilent 6890N gas chromatograph equipped with a (5%-phenyl)-methylpolysiloxane HP-5 column (length, 30 m; inside diameter, 0.32 mm; film thickness, 0.25 mm; Agilent Technologies, Ratingen, Germany) and a flame ionization detector (FID). Both heterologously produced sesquiterpenoid samples and standard calibration samples were diluted in ethyl acetate (Rodriguez et al., 2014). Temperatures of the injector and FID were set to 240°C and 250°C, respectively. Each sample had a volume of 1 μL and was injected splitless with helium as a carrier gas. The column temperature was sequentially changed: starting at 100°C for 5 min, ramped at 10°C min^−1^ to 180°C and then at 20°C min^−1^ to 300°C. The signals of heterologously produced (+)-valencene, which were absent in control samples, were confirmed by comparison of retention time to a commercial reference of (+)-valencene purchased (75056, Merck). To estimate the titre of α-cuprenene, the chemically similar reference compound β-caryophyllene was used (22075, Merck), because α-cuprenene is not commercially available to correlate putative α-cuprenene signals, which were absent in control samples, to compound amounts.

### Data availability

All data generated or analysed during this study are included in the manuscript and/or the Supplementary Files.

## Supporting information

Supplementary_Material Lee et al.

## Author contributions

JL, AL, KEJ and MF designed and planned the study. JL performed experiments. FH and AL supported GC-FID analysis. JL, FH, AL, KEJ and MF analysed the data. JL and MF designed and revised the manuscript. MF directed the project.

## Acknowledgements

We acknowledge lab members for discussion and comments on the manuscript. We are grateful to Drs. L. Olgeiser, F. Hennicke and T. Drepper for continuous support. K. Müntjes helped with microscopic analysis and provided a control strain expressing *gfp*. Dr. S. Jankowski generated Gfp-SKL and Pex3-Gfp strains for peroxisomal localisation studies. K. Hußnätter provided information on the promoter P_*rpl40*_. Dr. V. D. Urlacher and A. Kokorin supported the GC analysis. B. D. Anderson assisted the project within the framework of a RISE-DAAD scholarship.

## Supplementary Material

The Supplementary Material for this article can be found online at: https://www./xx.

