## Supplementary_Material Lee et al. for "*Ustilago maydis* serves as a novel production host for the synthesis of plant and fungal sesquiterpenoids"

Including Supplementary Figure S1-4 as well as Supplementary Table S1-S4

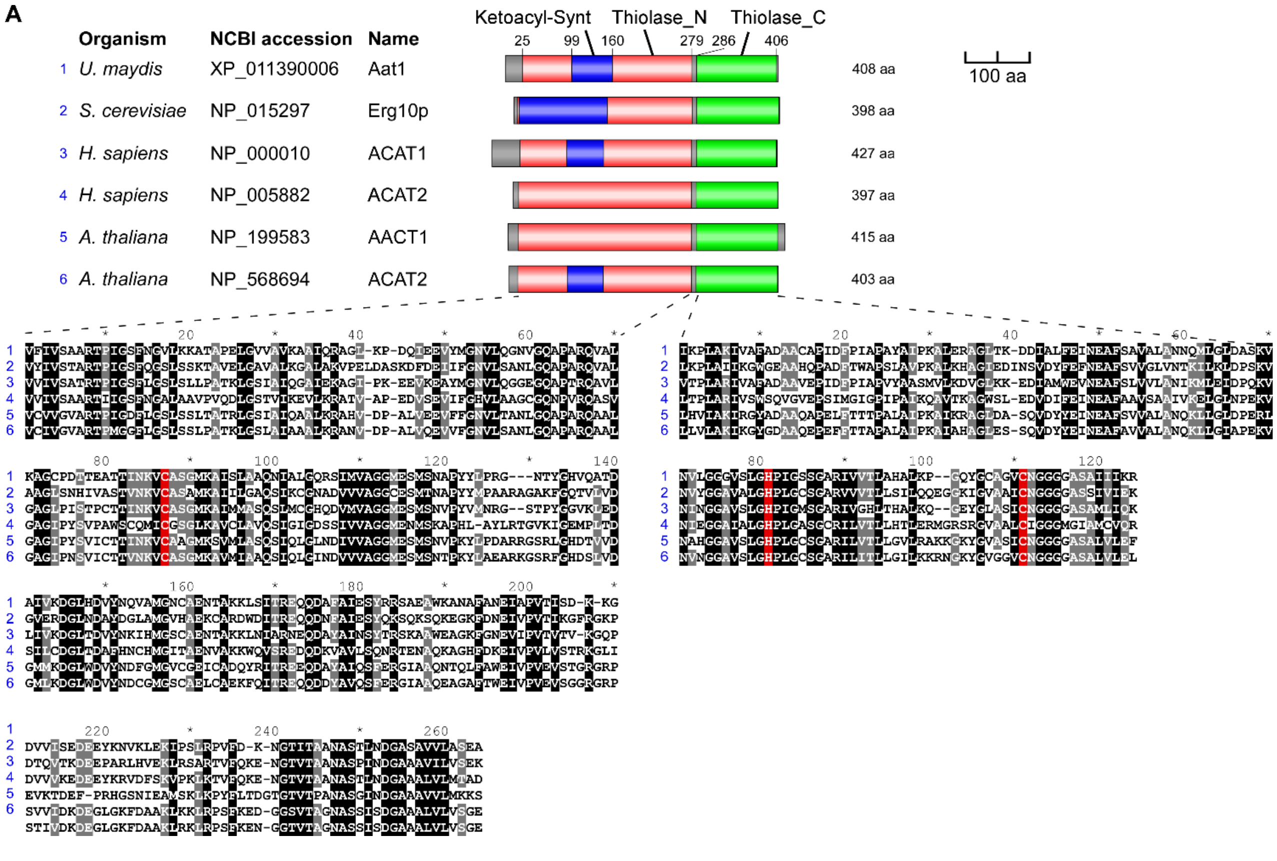


**Supplementary Figure S1. Bioinformatics analysis of *U. maydis* biosynthetic enzymes.**
(**A**) Amino acid sequence comparison of acetyl-CoA C-acetyltransferase Aat1 with homologues in fungi, animals and plants. Domain architecture according to SMART is given on the top (Ketoacyl-Synt, protein families Pfam identifier PF00109; Thiolase_N, PF00108; Thiolase_C, PF02803) and a sequence alignment of the corresponding domains is given at the bottom. Key amino acids in the active site of the enzyme from the bacterium *Zoogloea ramigera* are given in red (Miziorko, 2011).
(**B-H**) Continued on next page.

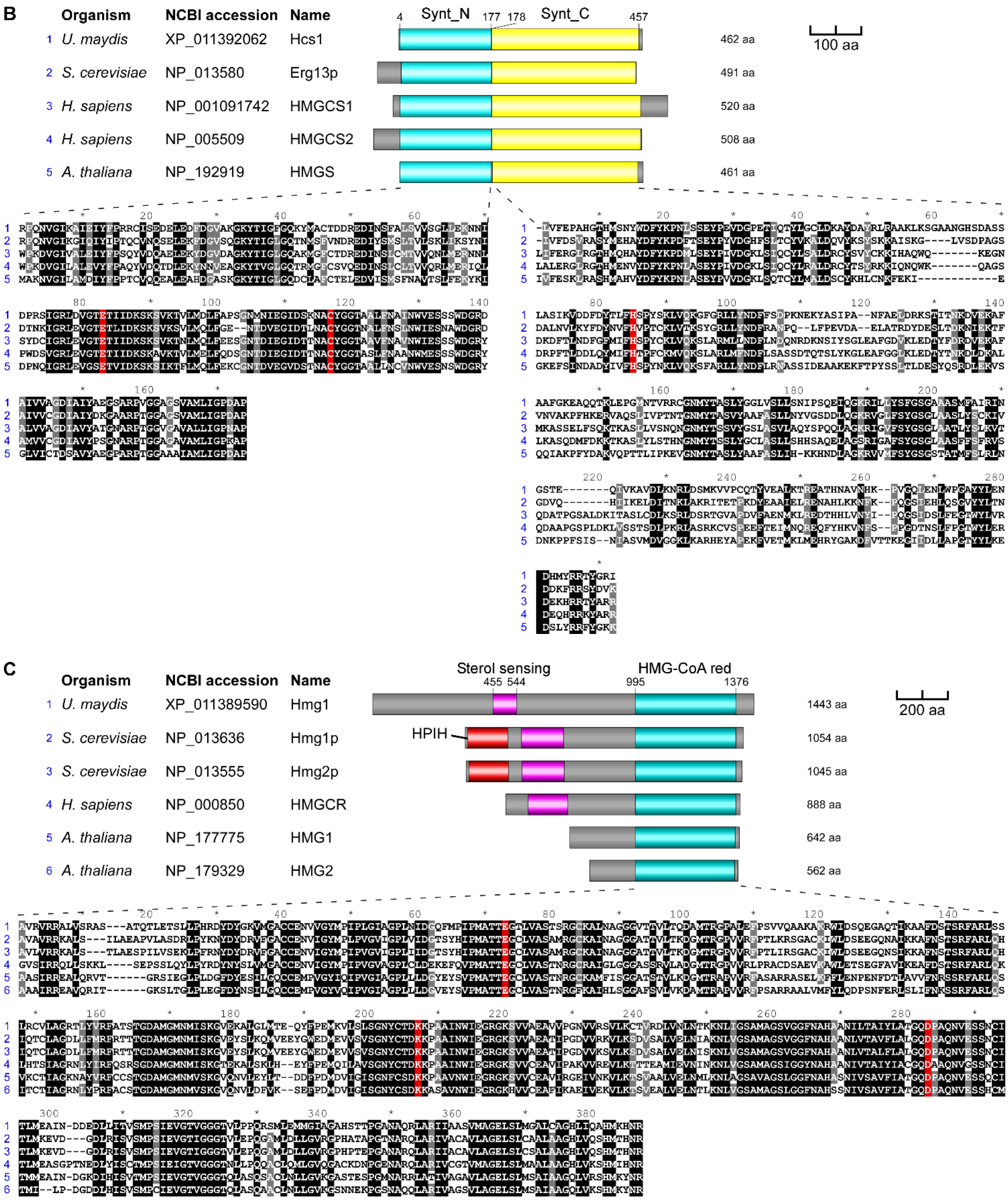


**Supplementary Figure S2. Bioinformatics analysis of *U. maydis* biosynthetic enzymes.** (**B-C**) Amino acid sequence comparison of 3-hydroxy-3-methylglutaryl-CoA synthases (B, active site from *Staphylococcus aureus*) and 3-hydroxy-3-methylglutaryl-CoA reductases (C, active site from *H. sapiens*) as shown in (A) (Miziorko, 2011; Synt_N, PF01154; Synt_C, PF08540; Sterol sensing, PF12349; HMG-CoA red, PF00368; HPIH, PF13323). (**D-H**) Continued on next page.

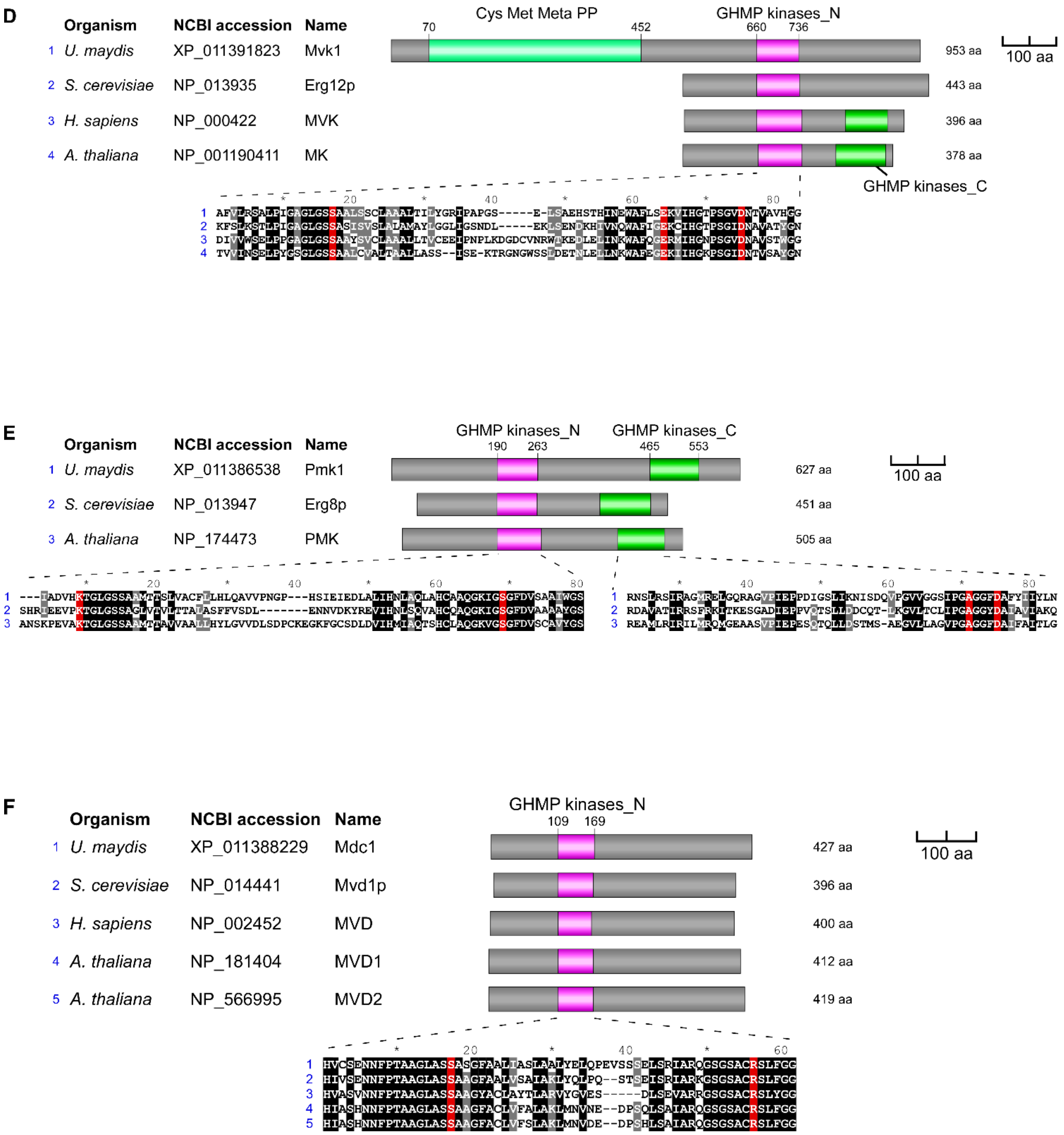


**Supplementary Figure S3. Bioinformatics analysis of *U. maydis* biosynthetic enzymes.**

(**D-F**) Amino acid sequence comparison of mevalonate kinases (D, active site from *Rattus norvegicus*) and phosphomevalonate kinases (E, active site from the bacterium *Streptococcus pneumoniae*, Andreassi et al. 2009 Biochemistry 48:6461) and mevalonate diphosphate decarboxylases (F, active site from *H. sapiens*) as shown in (A) (Miziorko, 2011; Cys Met Meta PP, PF01053; GHMP kinases_N, PF00288; GHMP kinases_C, PF08544). (**G-H)** Continued on next page.

G

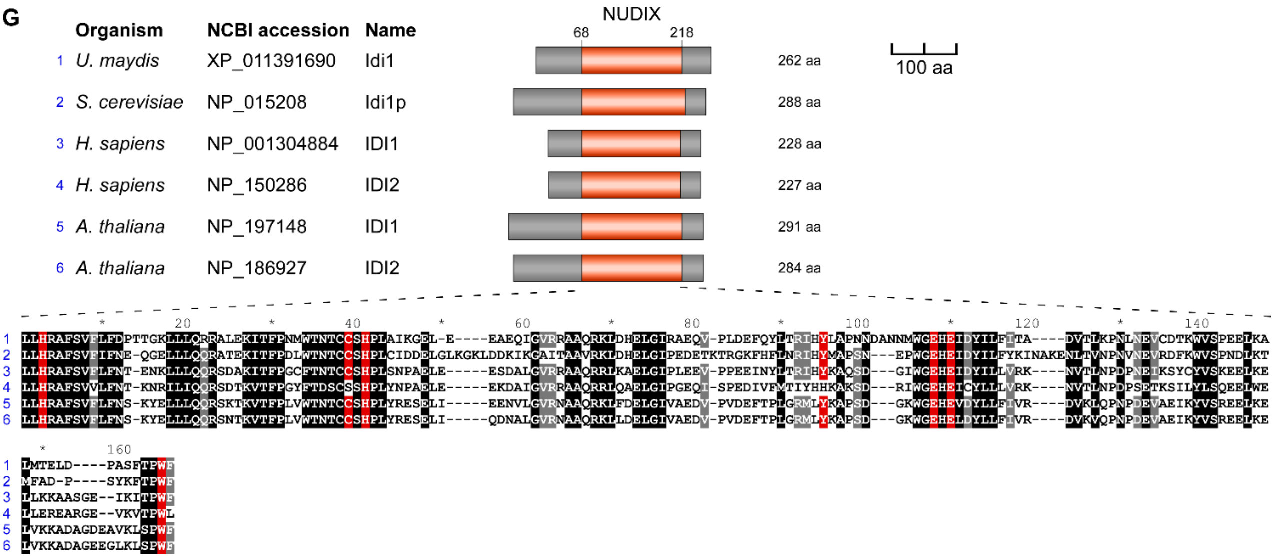


H

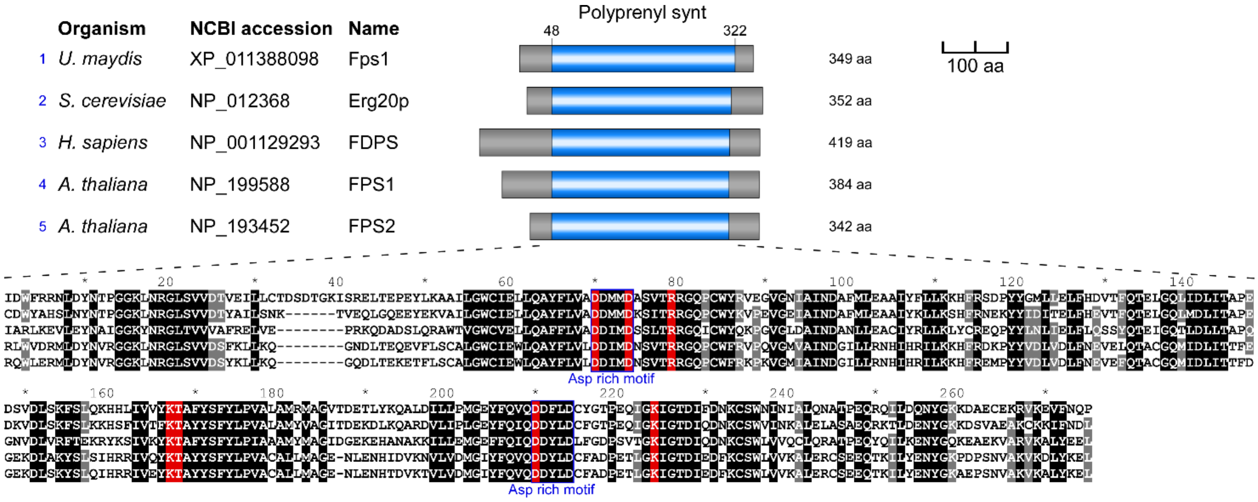


**Supplementary Figure S4. Bioinformatics analysis of *U. maydis* biosynthetic enzymes.**

(**G-H**) Amino acid sequence comparison of isopentenyl diphosphate isomerases (G, active site from *H. sapiens;* Zheng et al. 2007 *J. Mol. Biol.* 366:1447) and farnesyl diphosphate synthase/dimethylallyl transtransferases (H, active site from *H. sapiens*; Kavanagh et al. 2006 *PNAS* 103:7829) as shown in (A; NUDIX, PF00293; Polyprenyl synt, PF00348). End of figure.

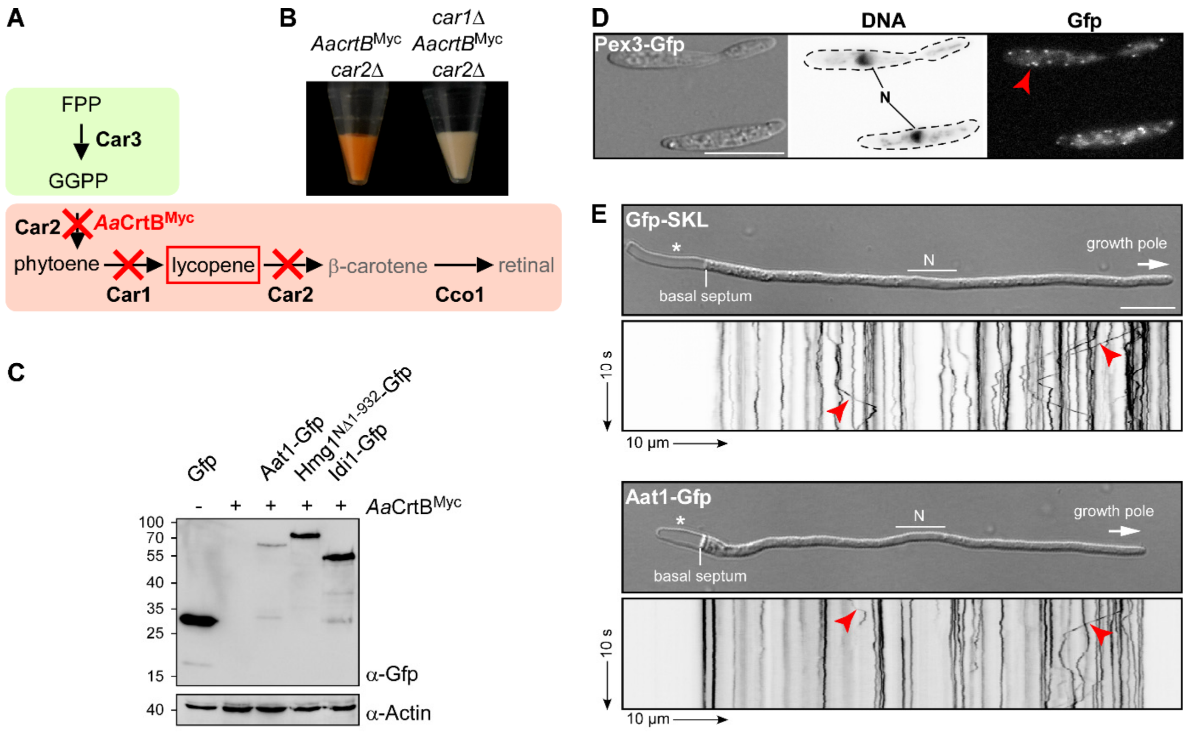


**Supplementary Figure S2. Aat1 localises to motile peroxisomes during hyphal growth.**

(**A**) Schematic representation of the carotenoid module given in Figure 1A (red cross indicates gene deletion). (**B**) Cell pellets of strains indicated above the image. (**C**) Western blot analysis of strains indicated above the lanes. The expected molecular weight is 70 kDa for Aat1-Gfp, 81 kDa for Hmg1$^{N\Delta 1\text{-}932}$-Gfp, 57 kDa for Idi1-Gfp, 37 kDa for *Aa*CrtB$^{Myc}$ and 57 kDa for actin (UMAG_11232). Antibodies are given in the lower right corner (size marker in kDa given on the left). Note, that the *Aa*CrtB$^{Myc}$ expressing strains carried a deletion of *car2*. (**D**) Microscopic analysis showing DIC image of fixed cells on the left (size bar, 10 µm). Corresponding staining of nuclear DNA with Hoechst 33342 (middle panel; N, nucleus, inverted image) and green fluorescence (Gfp) on the right (red arrowheads indicate peroxisomes). (**E**) Microscopic analysis showing DIC images of AB33 hyphae (6 h.p.i.) on top (size bar, 10 µm; N, nucleus) and corresponding kymographs at the bottom; genetic background as indicated (arrow length on the left and bottom indicates time and distance, respectively). Bidirectional movement of peroxisomes is visible as diagonal lines (red arrowheads).

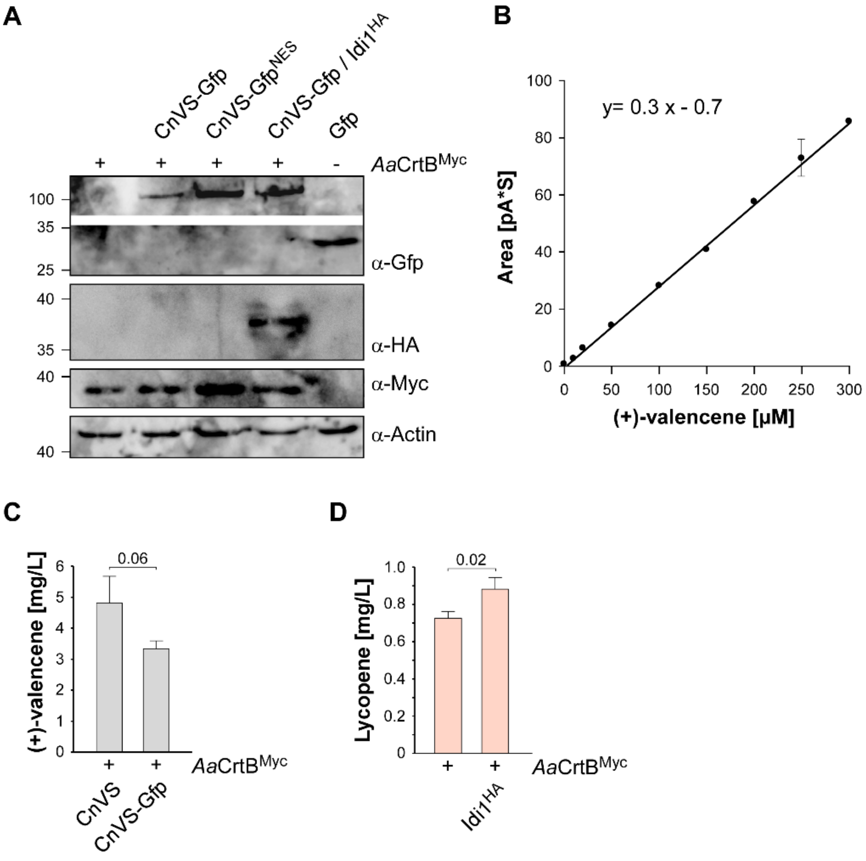


**Supplementary Figure S3. Expression of (+)-valencene synthase CnVS in *U. maydis.***

(**A**) Western blot analysis of strains indicated above the lanes. The expected molecular weight is 96 kDa for CnVS-Gfp, 97 kDa for CnVS-Gfp$^{NES}$, 27 kDa for Gfp, 37 kDa for *Aa*CrtB$^{Myc}$ and 42 kDa for actin (UMAG_11232). Antibodies are given in the lower right corner (size marker in kDa given on the left). Note, that the *Aa*CrtB$^{Myc}$ expressing strain carried a deletion of *car2*. (**B**) Standard calibration curve of the peaks derived from GC-FID measurements versus (+)-valencene concentration. (**C**) Concentration of (+)-valencene in the n-dodecane samples were determined based on the calibration curve with the commercial reference compound used in (B). Three independent biological experiments (n=3) were carried out. Error bars indicate standard deviation of the mean (SD). Statistical significance was calculated using the unpaired two-tailed *t* test and *p*-values were indicated above. Note, that the *Aa*CrtB$^{Myc}$ expressing strains carried a deletion of *car2*. (**D**) Lycopene concentrations in strains given at the bottom. Three independent biological experiments (n=3) were carried out. Error bars indicate standard deviation of the mean (SD). Statistical significance was calculated using the unpaired two-tailed *t* test and *p*-values were indicated above. Note, that the *Aa*CrtB$^{Myc}$ expressing strains carried a deletion of *car2*.

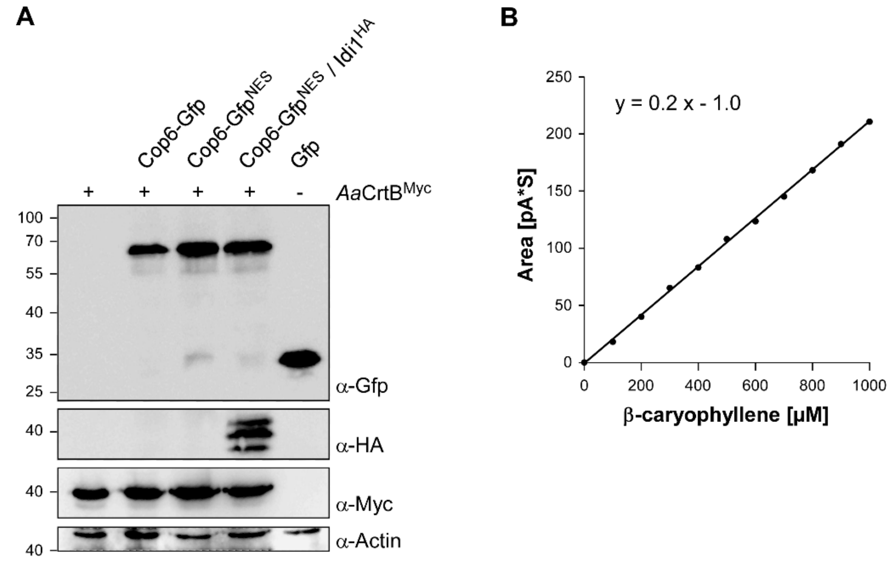


**Supplementary Figure S4. Heterologous expression of Cop6 in *U. maydis*.**

(**A**) Western blot analysis of strains indicated above the lanes. The expected molecular weight is 65 kDa for Cop6-Gfp, 66 kDa for Cop6-Gfp$^{NES}$, 27 kDa for Gfp, 37 kDa for *Aa*CrtB$^{Myc}$ and 42 kDa for actin (UMAG_11232). Antibodies are given in the lower right corner (size marker in kDa given on the left). Note, that the *Aa*CrtB$^{Myc}$ expressing strains carried a deletion of *car2*. (**B**) Standard calibration curve of the peaks derived from GC-FID measurements versus the chemically similar compound β-caryophyllene.

### Supplementary Table S1: Description of *U. maydis* strains used in this study

| Strain | Locus | Progenitor strain | Short description |
|---|---|---|---|
| AB33 | *b* | FB2 | *Pnar:bW2bE1*, expression of active b heterodimer under control of the $P_{nar1}$ promoter; strain grows filamentously upon changing the nitrogen source. Published in (Brachmann, 2001). |
| AB33car2Δ_HygR | *car2* | AB33 | Carrying a deletion of *car2* and possessing hygromycin B resistance. |
| AB33car2Δ/AacrtB$^{Myc}$_NatR | *car2* | AB33car2Δ_HygR | Carrying a deletion of *car2* and expressing codon-optimized *Aa*CrtB, which is C-terminally fused to 3× Myc tag at the *car2* locus. Possessing nourseothricin resistance. |
| AB33car2Δ/AacrtB$^{Myc}$_NatR/ car1Δ_HygR | *car2* *car1* | AB33car2Δ/AacrtB$^{Myc}$_NatR | Carrying deletions of *car2* and *car1* and expressing codon-optimized *Aa*CrtB, which is C-terminally fused to 3× Myc tag at the *car2* locus. Possessing nourseothricin and hygromycin B resistance. |
| AB33car2Δ/AacrtB$^{Myc}$_NatR/ aat1$^{HA}$_CbxR | *car2* *ip$^s$* | AB33car2Δ/AacrtB$^{Myc}$_NatR | Carrying a deletion of *car2* and co-expressing codon-optimized *Aa*CrtB, which is C-terminally fused to 3× Myc tag at the *car2* locus, and Aat1 N-terminally fused to 3× HA tag at the *ip$^s$* locus. Possessing nourseothricin and carboxin resistance. |
| AB33car2Δ/AacrtB$^{Myc}$_NatR/ hmg1$^{NΔ1\text{-}932HA}$_CbxR | *car2* *ip$^s$* | AB33car2Δ/AacrtB$^{Myc}$_NatR | Carrying a deletion of *car2* and co-expressing codon-optimized *Aa*CrtB, which is C-terminally fused to 3× Myc tag at the *car2* locus, and N-terminally truncated from aa 1 to 932 version of Hmg1, which is N-terminally fused to 3× HA tag at the *ip$^s$* locus. Possessing nourseothricin and carboxin resistance. |
| AB33car2Δ/AacrtB$^{Myc}$_NatR/ idi1$^{HA}$_CbxR | *car2* *ip$^s$* | AB33car2Δ/AacrtB$^{Myc}$_NatR | Carrying a deletion of *car2* and co-expressing codon-optimized *Aa*CrtB, which is C-terminally fused to 3× Myc tag at the *car2* locus, and Idi1 N-terminally fused to 3× HA tag at the *ip$^s$* locus. possessing nourseothricin and carboxin resistance. |
| AB33car2Δ/AacrtB$^{Myc}$_NatR/ aat1G_CbxR | *car2* *ip$^s$* | AB33car2Δ/AacrtB$^{Myc}$_NatR | Carrying a deletion of *car2* and co-expressing codon-optimized *Aa*CrtB, which is C-terminally fused to 3× Myc tag at the *car2* locus, and Aat1 N-terminally fused to eGfp at the *ip$^s$* locus. Possessing nourseothricin and carboxin resistance. |
| AB33car2Δ/AacrtB$^{Myc}$_NatR/ hmg1$^{NΔ1\text{-}932}$G_CbxR | *car2* *ip$^s$* | AB33car2Δ/AacrtB$^{Myc}$_NatR | Carrying a deletion of *car2* and co-expressing codon-optimized *Aa*CrtB, which is C-terminally fused to 3× Myc tag at the *car2* locus, and N-terminally truncated from aa 1 to 932 version of Hmg1, which is N-terminally fused to eGfp at the *ip$^s$* locus. Possessing nourseothricin and carboxin resistance. |
| AB33car2Δ/AacrtB$^{Myc}$_NatR/ idi1G_CbxR | *car2* *ip$^s$* | AB33car2Δ/AacrtB$^{Myc}$_NatR | Carrying a deletion of *car2* and co-expressing codon-optimized *Aa*CrtB, which is C-terminally fused to 3× Myc tag at the *car2* locus and Idi1 N-terminally fused to eGfp at the *ip$^s$* locus. Possessing nourseothricin and carboxin resistance. |
| AB33egfp_CbxR | *ip$^s$* | AB33 | The *egfp* construct is ectopically integrated into the *ip$^s$* locus. Possessing nourseothricin and carboxin resistance. Published in (Köpke, 2011). |
| AB33upa9mC_HygR/ egfp-skl_CbxR | *upa9* *ip$^s$* | AB33egfp-skl_CbxR | Co-expressing eGfp containing SKL at the C-terminus at the *ip$^s$* locus and Upa9 C-terminally fused to eGfp at the *upa9* locus. Upa9 is the homolog of Pex3. |
| AB33pab1mC_HygR/ upa9G_NatR | *pab1* *upa9* | AB33pab1mC | Co-expressing Pab1 C-terminally fused to mCherry at the *pab1* locus and Upa9 C-terminally fused to eGfp at the *upa9* locus. Possessing hygromycin B and carboxin resistance. |
| AB33car2Δ/AacrtB$^{Myc}$ | *car2* | AB33car2Δ/AacrtB$^{Myc}$_NatR | Carrying a deletion of *car2* and expressing codon-optimized *Aa*CrtB, which is C-terminally fused to 3× Myc tag at the *car2* locus. Nourseothricin resistance gene cassette is recycled via FRTm7 |
| AB33car2Δ/AacrtB$^{Myc}$/ cco1Δ_HygR | *car2* *cco1* | AB33car2Δ/AacrtB$^{Myc}$ | Carrying deletions of *car2* and *cco1* and expressing codon-optimized *Aa*CrtB, which is C-terminally fused to 3× Myc tag at the *car2* locus. Nourseothricin resistance gene cassette is recycled via FRTm7 and possessing hygromycin B resistance. |
| AB33car2Δ/AacrtB$^{Myc}$/cco1Δ/ idi1$^{HA}$_NatR | *car2* *cco1* | AB33car2Δ/AacrtB$^{Myc}$/cco1Δ _HygR | Carrying deletions of *car2* and *cco1* and co-expressing codon-optimized *Aa*CrtB, which is C-terminally fused to 3× Myc tag at the *car2* locus, and Idi1 N-terminally fused to 3× HA tag at the *cco1* locus. Possessing nourseothricin resistance. |

| | | | |
|---|---|---|---|
| AB33car2Δ/AacrtB$^{Myc}$/cco1Δ/ idi1$^{HA}$ | *car2* *cco1* | AB33car2Δ/AacrtB$^{Myc}$/cco1Δ /idi1$^{HA}$_NatR | Carrying deletions of *car2* and *cco1* and co-expressing codon-optimized *Aa*CrtB, which is C-terminally fused to 3× Myc tag at the *car2* locus, and Idi1 N-terminally fused to 3× HA tag at the *cco1* locus. Nourseothricin resistance gene cassette is recycled via FRTm1. |
| AB33car2Δ/AacrtB$^{Myc}$/cco1Δ/ idi1$^{HA}$/upp3Δ_HygR | *car2* *cco1* | AB33car2Δ/AacrtB$^{Myc}$/cco1Δ /idi1$^{HA}$ | Carrying deletions of *car2*, *cco1*, and *upp3*, and co-expressing codon-optimized *Aa*CrtB, which is C-terminally fused to 3× Myc tag at the *car2* locus, and Idi1 N-terminally fused to 3× HA tag at the *cco1* locus. Possessing hygromycin B resistance. |
| AB33upp3Δ/egfp_NatR | *upp3* | AB33upp3Δ_HygR | Carrying a deletion of *upp3* and expressing eGfp at the *upp3* locus. Possessing nourseothricin resistance. |
| AB33car2Δ/AacrtB$^{Myc}$/ upp3Δ_HygR | *car2* *upp3* | AB33car2Δ/AacrtB$^{Myc}$ | Carrying deletions of *car2* and *upp3* and expressing codon-optimized *Aa*CrtB, which is C-terminally fused to 3× Myc tag at the *car2* locus. Possessing hygromycin B resistance |
| AB33car2Δ/AacrtB$^{Myc}$/upp3Δ/ CnVS_NatR | *car2* *upp3* | AB33car2Δ/AacrtB$^{Myc}$/upp3Δ _HygR | Carrying deletions of *car2* and *upp3* and co-expressing codon-optimized *Aa*CrtB, which is C-terminally fused to 3× Myc tag at the *car2* locus, and dicodon-optimized CnVS at the *upp3* locus. Possessing nourseothricin resistance. |
| AB33car2Δ/AacrtB$^{Myc}$/upp3Δ/ CnVSG_NatR | *car2* *upp3* | AB33car2Δ/AacrtB$^{Myc}$/upp3Δ _HygR | Carrying deletions of *car2* and *upp3* and co-expressing codon-optimized *Aa*CrtB, which is C-terminally fused to 3× Myc tag at the *car2* locus, and dicodon-optimized CnVS, which is N-terminally fused to eGfp at the *upp3* locus. Possessing nourseothricin resistance. |
| AB33car2Δ/AacrtB$^{Myc}$/upp3Δ/ CnVSG$^{NES}$_NatR | *car2* *upp3* | AB33car2Δ/AacrtB$^{Myc}$/upp3Δ _HygR | Carrying deletions of *car2* and *upp3* and co-expressing codon-optimized *Aa*CrtB, which is C-terminally fused to 3× Myc tag at the *car2* locus, and dicodon-optimized CnVS, which is N-terminally fused to eGfp$^{NES}$ at the *upp3* locus. Possessing nourseothricin resistance. |
| AB33car2Δ/AacrtB$^{Myc}$/cco1Δ/ idi1$^{HA}$/upp3Δ/CnVSG$^{NES}$_NatR | *car2* *cco1* *upp3* | AB33car2Δ/AacrtB$^{Myc}$/cco1Δ /idi1$^{HA}$/upp3Δ_HygR | Carrying deletions of *car2*, *cco1*, and *upp3* and co-expressing codon-optimized *Aa*CrtB, which is C-terminally fused to 3× Myc tag at the *car2* locus, Idi1 N-terminally fused to 3× HA tag at the *cco1* locus, and dicodon-optimized CnVS, which is N-terminally fused to eGfp$^{NES}$ at the *upp3* locus. Possessing nourseothricin resistance. |
| AB33car2Δ/AacrtB$^{Myc}$/upp3Δ/ cop6_NatR | *car2* *upp3* | AB33car2Δ/AacrtB$^{Myc}$/upp3Δ _HygR | Carrying deletions of *car2* and *upp3* and co-expressing codon-optimized *Aa*CrtB, which is C-terminally fused to 3× Myc tag at the *car2* locus, and dicodon-optimized Cop6 at the *upp3* locus. Possessing nourseothricin resistance. |
| AB33car2Δ/AacrtB$^{Myc}$/upp3Δ/ cop6G_NatR | *car2* *upp3* | AB33car2Δ/AacrtB$^{Myc}$/upp3Δ _HygR | Carrying deletions of *car2* and *upp3* and co-expressing codon-optimized *Aa*CrtB, which is C-terminally fused to 3× Myc tag at the *car2* locus, and dicodon-optimized Cop6, which is N-terminally fused to eGfp at the *upp3* locus. Possessing nourseothricin resistance. |
| AB33car2Δ/AacrtB$^{Myc}$/upp3Δ/ cop6G$^{NES}$_NatR | *car2* *upp3* | AB33car2Δ/AacrtB$^{Myc}$/upp3Δ _HygR | Carrying deletions of *car2* and *upp3* and co-expressing codon-optimized *Aa*CrtB, which is C-terminally fused to 3× Myc tag at the *car2* locus and dicodon-optimized Cop6, which is N-terminally fused to eGfp$^{NES}$ at the *upp3* locus. Possessing nourseothricin resistance. |
| AB33car2Δ/AacrtB$^{Myc}$/cco1Δ/ idi1$^{HA}$/upp3Δ/cop6G$^{NES}$_NatR | *car2* *cco1* *upp3* | AB33car2Δ/AacrtB$^{Myc}$/cco1Δ /idi1$^{HA}$/upp3Δ_HygR | Carrying deletions of *car2*, *cco1*, and *upp3* and co-expressing codon-optimized *Aa*CrtB, which is C-terminally fused to 3× Myc tag at the *car2* locus, Idi1 N-terminally fused to 3× HA tag at the *cco1* locus, and dicodon-optimized Cop6, which is N-terminally fused to eGfp$^{NES}$ at the *upp3* locus. Possessing nourseothricin resistance. |

### Supplementary Table S2: Generation of *U. maydis* strains used in this study

| Strain | Relevant genotype | UMa | Reference | Transformed plasmid (pUMa) | Locus | Progenitor strain |
|---|---|---|---|---|---|---|
| AB33 | *a2 Pnar::bW2bE1* | 133 | Brachmann, 2001 | pAB33 | *b* | FB2 |
| AB33car2Δ_HygR | *car2Δ* | 2290 | This study | pCar2Δ_HygR$^{FRT}$ (pUMa3287) | *car2* | AB33 |
| AB33car2Δ/ AacrtB$^{Myc}$_NatR | *car2Δ/AacrtB* | 2612 | This study | pAacrtB$^{Myc}$_NatR$^{FRTm7}$ (pUMa3707) | *car2* | AB33car2Δ_HygR |
| AB33car2Δ/AacrtB$^{Myc}$_NatR/ car1Δ_HygR | *car2Δ/AacrtB/ car1Δ* | 3246 | This study | pCar1Δ_HygR$^{FRT}$ (pUMa3286) | *car1* | AB33car2Δ/ AacrtB$^{Myc}$_NatR |
| AB33car2Δ/AacrtB$^{Myc}$_NatR/ aat1$^{HA}$_CbxR | *car2Δ/AacrtB/ aat1* | 2706 | This study | pAat1$^{HA}$_CbxR (pUMa3379) | *ip$^{s}$* | AB33car2Δ/ AacrtB$^{Myc}$_NatR |
| AB33car2Δ/AacrtB$^{Myc}$_NatR/ hmg1$^{NΔ1-932HA}$_CbxR | *car2Δ/AacrtB/ hmg1$^{NΔ1-932}$* | 2707 | This study | pHmg1$^{NΔ1-932}$_CbxR (pUMa3381) | *ip$^{s}$* | AB33car2Δ/ AacrtB$^{Myc}$_NatR |
| AB33car2Δ/AacrtB$^{Myc}$_NatR/ idi1$^{HA}$_CbxR | *car2Δ/AacrtB/ idi1* | 2708 | This study | pIdi1$^{HA}$_CbxR (pUMa3382) | *ip$^{s}$* | AB33car2Δ/ AacrtB$^{Myc}$_NatR |
| AB33car2Δ/AacrtB$^{Myc}$_NatR/ aat1G_CbxR | *car2Δ/AacrtB/ aat1G* | 3065 | This study | pAat1G_CbxR (pUMa4230) | *ip$^{s}$* | AB33car2Δ/ AacrtB$^{Myc}$_NatR |
| AB33car2Δ/AacrtB$^{Myc}$_NatR/ hmg1$^{NΔ1-932}$G_CbxR | *car2Δ/AacrtB/ hmg1$^{NΔ1-932}$* | 3066 | This study | pHmg1$^{NΔ1-932}$G_CbxR (pUMa4231) | *ip$^{s}$* | AB33car2Δ/ AacrtB$^{Myc}$_NatR |
| AB33car2Δ/AacrtB$^{Myc}$_NatR/ idi1G_CbxR | *car2Δ/AacrtB/ idi1G* | 3067 | This study | pIdi1G_CbxR (pUMa4232) | *ip$^{s}$* | AB33car2Δ/ AacrtB$^{Myc}$_NatR |
| AB33egfp_CbxR | *egfp* | 2229 | Köpke, 2011 | peGfp_CbxR (pUMa1139) | *ip$^{s}$* | AB33 |
| AB33upa9mC_HygR/ egfp-skl_CbxR | *upa9mC/egfp-skl* | 2386 | This study | pUpa9mC_HygR (pUMa2722) | *upa9* | AB33egfp-skl_CbxR |
| AB33pab1mC_HygR/ upa9G_NatR | *pab1mC/upa9G* | 1951 | This study | pUpa9G_NatR (pUMa2938) | *upa9* | AB33pab1mC |
| AB33car2Δ/AacrtB$^{Myc}$ | *car2Δ/AacrtB* | 2775 | This study | pFLPexpC (pUMa1446) | *car2* | AB33car2Δ/ AacrtB$^{Myc}$_NatR |
| AB33car2Δ/AacrtB$^{Myc}$/ cco1Δ_HygR | *car2Δ/AacrtB/ cco1Δ* | 2785 | This study | pCco1Δ_HygR (pUMa3281) | *cco1* | AB33car2Δ/ AacrtB$^{Myc}$ |
| AB33car2Δ/AacrtB$^{Myc}$/ cco1Δ/idi1$^{HA}$_NatR | *car2Δ/AacrtB/ cco1Δ/idi1* | 3025 | This study | pIdi1$^{HA}$_NatR$^{FRTm1}$ (pUMa4082) | *cco1* | AB33car2Δ/AacrtB$^{Myc}$/ cco1Δ_HygR |
| AB33car2Δ/AacrtB$^{Myc}$/ cco1Δ/idi1$^{HA}$ | *car2Δ/AacrtB/ cco1Δ/idi1* | 3077 | This study | pFLPexpC (pUMa1446) | *cco1* | AB33car2Δ/AacrtB$^{Myc}$/ cco1Δ/idi1$^{HA}$_NatR |
| AB33car2Δ/AacrtB$^{Myc}$/ cco1Δ/idi1$^{HA}$/upp3Δ_HygR | *car2Δ/AacrtB/ cco1Δ/idi1/upp3Δ* | 3131 | This study | pUpp3Δ_HygR$^{FRTm3}$ (pUMa1556) | *upp3* | AB33car2Δ/AacrtB$^{Myc}$/ cco1Δ/idi1$^{HA}$ |
| AB33upp3Δ/egfp_NatR | *upp3Δ/egfp* | 2179 | This study | peGfp_NatR (pUMa3132) | *upp3* | AB33upp3Δ_HygR |
| AB33car2Δ/AacrtB$^{Myc}$/ upp3Δ_HygR | *car2Δ/AacrtB/ upp3Δ* | 2884 | This study | pUpp3Δ_HygR$^{FRTm3}$ (pUMa1556) | *upp3* | AB33car2Δ/AacrtB$^{Myc}$ |
| AB33car2Δ/AacrtB$^{Myc}$/ upp3Δ/CnVS_NatR | *car2Δ/AacrtB/ upp3Δ/CnVS* | 3194 | This study | pCnVS_NatR$^{FRTm2}$ (pUMa4500) | *upp3* | AB33car2Δ/AacrtB$^{Myc}$/ upp3Δ_HygR |
| AB33car2Δ/AacrtB$^{Myc}$/ upp3Δ/CnVSG_NatR | *car2Δ/AacrtB/ upp3Δ/CnVSG* | 3104 | This study | pCnVSG_NatR$^{FRTm2}$ (pUMa4356) | *upp3* | AB33car2Δ/AacrtB$^{Myc}$/ upp3Δ_HygR |
| AB33car2Δ/AacrtB$^{Myc}$/ upp3Δ/CnVSG$^{NES}$_NatR | *car2Δ/AacrtB/ upp3Δ/CnVSG$^{NES}$* | 3192 | This study | pCnVSG$^{NES}$_NatR$^{FRTm2}$ (pUMa4499) | *upp3* | AB33car2Δ/AacrtB$^{Myc}$/ upp3Δ_HygR |
| AB33car2Δ/AacrtB$^{Myc}$/ cco1Δ/idi1$^{HA}$/ upp3Δ/CnVSG$^{NES}$_NatR | *car2Δ/AacrtB/ cco1Δ/idi1/ upp3Δ/CnVSG$^{NES}$* | 3195 | This study | pCnVSG$^{NES}$_NatR$^{FRTm2}$ (pUMa4499) | *upp3* | AB33car2Δ/AacrtB$^{Myc}$/ cco1Δ/idi1$^{HA}$/ upp3Δ_HygR |
| AB33car2Δ/AacrtB$^{Myc}$/ upp3Δ/cop6_NatR | *car2Δ/AacrtB/ upp3Δ/cop6* | 3193 | This study | pCop6_NatR$^{FRTm2}$ (pUMa4496) | *upp3* | AB33car2Δ/AacrtB$^{Myc}$/ upp3Δ_HygR |
| AB33car2Δ/AacrtB$^{Myc}$/ upp3Δ/cop6G_NatR | *car2Δ/AacrtB/ upp3Δ/cop6G* | 2944 | This study | pCop6G_NatR$^{FRTm2}$ (pUMa4089) | *upp3* | AB33car2Δ/AacrtB$^{Myc}$/ upp3Δ_HygR |
| AB33car2Δ/AacrtB$^{Myc}$/ upp3Δ/cop6G$^{NES}$_NatR | *car2Δ/AacrtB/ upp3Δ/cop6G$^{NES}$* | 3078 | This study | pCop6G$^{NES}$_NatR$^{FRTm2}$ (pUMa4316) | *upp3* | AB33car2Δ/AacrtB$^{Myc}$/ upp3Δ_HygR |
| AB33car2Δ/AacrtB$^{Myc}$/ cco1Δ/idi1$^{HA}$/ upp3Δ/cop6G$^{NES}$_NatR | *car2Δ/AacrtB/ cco1Δ/idi1/ upp3Δ/cop6G$^{NES}$* | 3148 | This study | pCop6G$^{NES}$_NatR$^{FRTm2}$ (pUMa4316) | *upp3* | AB33car2Δ/AacrtB$^{Myc}$/ cco1Δ/idi1$^{HA}$/ upp3Δ_HygR |

### Supplementary Table S3: Description of plasmids used for *U. maydis* strains generation

| Plasmid name | pUMa | Resistance cassette | Short description |
|---|---|---|---|
| pCco1Δ_HygR | 3281 | *Sfi*I-insert MF1hs | Plasmid for generating deletion mutants of *cco1*. The hygromycin B resistance cassette contains FRT sites (GAAGTTCCTATTCTCTAGAAA GTATAGGAACTTC) at both ends for recycling by Flp recombinase. The cassette is flanked by the regions, 1.1 kb upstream and 0.8 kb downstream of *cco1*. The flanking regions were amplified by PCR using oMB981/oMB982 and oMB983/oMB984 and UM521 wild type DNA as template. |
| pCar1Δ_HygR | 3286 | *Sfi*I-insert MF1hs | Plasmid for generating deletion mutants of *car1*. The hygromycin B resistance cassette contains FRT sites at both ends for recycling by Flp recombinase. The cassette is flanked by the regions, 0.9 kb upstream and 0.6 kb downstream of *car1*. The flanking regions were amplified by PCR using oUP134/oMB998 and oMB999/oUP009 and UM521 wild type DNA as template. |
| pCar2Δ_HygR | 3287 | *Sfi*I-insert MF1hs | Plasmid for generating deletion mutants of *car2*. The hygromycin B resistance cassette contains FRT sites at both ends for recycling by Flp recombinase. The cassette is flanked by the regions, 0.9 kb upstream and 0.9 kb downstream of *car2*. The flanking regions were amplified by PCR using oUP014/oUP015 and oUP016/oUP017 and UM521 wild type DNA as template. |
| pUpp3Δ_HygR | 1556 | *Sfi*I-insert MF1hs | Plasmid for generating deletion mutants of *upp3*. The hygromycin B resistance cassette contains FRTm3 sites (GAAGTTCCTATTCTCCAGA AAGTATAGGAACTTC) at both ends for recycling by Flp recombinase. The cassette is flanked by the regions, 1.5 kb upstream and 1.9 kb downstream of *upp3*. The flanking regions were amplified by PCR using UM521 wild type as template. Published (Sarkari, 2014). |
| pFLPexpC | 1446 | CbxR | Plasmid for expressing Flp recombinase, which is under the control of $P_{crg1}$ promoter and the expression is induced in the presence of arabinose. Containing the carboxin resistance cassette. Published (Khrunyk, 2010) |
| peGfp_NatR | 3132 | *Sfi*I-insert of pMF5-1n | Plasmid for expressing eGfp with the nourseothricin resistance cassette at the *upp3* locus. The expression of *egfp* is under the control of $P_{otef}$ and $T_{nos}$ terminator. |
| pAacrtB$^{Myc}$_NatR$^{FRTm7}$ | 3707 | *Sfi*I-insert of pMF5-1n | Plasmid for expressing codon-optimized *Aa*CrtB, which is C-terminally fused to 3× Myc tag with the nourseothricin resistance cassette containing FRTm7 (GAAGTTCCTATTCTCTATAAAGTATAGGAACTTC), at the *car2* locus. The expression of *AacrtB*$^{Myc}$ is under the control of $P_{rpl40}$ (1 kb upstream of the ORF) promoter and $T_{nos}$ terminator. |
| pAat1$^{HA}$_CbxR | 3379 | CbxR for integration at the *ip*$^{s}$ locus | Plasmid for expressing Aat1 N-terminally fused to 3× HA tag at the *ip*$^{s}$ locus. The expression is under the control of $P_{otef}$ promoter and $T_{nos}$ terminator. |
| pHmg1$^{N\Delta 1\text{-}932HA}$_CbxR | 3381 | CbxR for integration at the *ip*$^{s}$ locus | Plasmid for expressing N-terminally truncated from aa 1 to 932 of Hmg1, which is N-terminally fused to 3× HA tag, at the *ip*$^{s}$ locus. The expression is under the control of $P_{otef}$ promoter and $T_{nos}$ terminator. |
| pIdi1$^{HA}$_CbxR | 3382 | CbxR for integration at the *ip*$^{s}$ locus | Plasmid for expressing Idi1 N-terminally fused to 3× HA tag at the *ip*$^{s}$ locus. The expression is under the control of $P_{otef}$ promoter and $T_{nos}$ terminator. |
| pAat1G_CbxR | 4230 | CbxR for integration at the *ip*$^{s}$ locus | Plasmid for expressing Aat1 N-terminally fused to eGfp at the *ip*$^{s}$ locus. The expression is under the control of $P_{tef}$ promoter and $T_{nos}$ terminator. |
| pHmg1$^{N\Delta 1\text{-}932}$G_CbxR | 4231 | CbxR for integration at the *ip*$^{s}$ locus | Plasmid for expressing N-terminally truncated from aa 1 to 932 of Hmg1, which is N-terminally fused to eGfp, at the *ip*$^{s}$ locus. The expression is under the control of $P_{tef}$ promoter and $T_{nos}$ terminator. |
| pIdi1G_CbxR | 4232 | CbxR for integration at the *ip*$^{s}$ locus | Plasmid for expressing Idi1 N-terminally fused to eGfp at the *ip*$^{s}$ locus. The expression is under the control of $P_{tef}$ promoter and $T_{nos}$ terminator. |
| pUpa9mC_HygR | 2722 | *Sfi*I-insert MF1hs | Plasmid for expressing Upa9 C-terminally fused to mCherry with the hygromycin B resistance cassette at the *upa9* locus. The insert construct is flanked by the regions, 1.1 kb of *upa9* (UMAG_06200) ORF and 1.0 kb downstream of *upa9*. The flanking regions were amplified by PCR using oDD527/oDD605 and oDD634/oDD635 and UM521 wild type DNA as template. The expression is under the control of the native promoter and $T_{nos}$ terminator. |
| peGfp-skl_CbxR | 3141 | CbxR for integration at the *ip*$^{s}$ locus | Plasmid for expressing eGfp C-terminally fused to SKL at the *ip*$^{s}$ locus. The expression is under the control of $P_{otef}$ promoter and $T_{nos}$ terminator. |
| pUpa9G_NatR | 2938 | *Sfi*I-insert of pMF5-1n | Plasmid for expressing Upa9 C-terminally fused to eGfp with the nourseothricin resistance cassette at the *upa9* locus. The insert construct is flanked by the regions, 1.1 kb of *upa9* ORF and 1.0 kb downstream of *upa9*. The flanking regions were amplified by PCR using oDD527/oDD605 and oDD634/oDD635 and UM521 wild type DNA as template. The expression is under the control the native promoter and $T_{nos}$ terminator. |

| | | | |
|---|---|---|---|
| pIdi1$^{HA}$_NatR$^{FRTm1}$ | 4082 | *Sfi*I-insert of pMF5-1n | Plasmid for expressing Idi1 N-terminally fused to 3× HA tag with the nourseothricin resistance cassette containing FRTm1 sites (GAAGTTCC TATTCTCGAGAAAGTATAGGAACTTC) at both ends for recycling by Flp recombinase at the *cco1* locus. The expression is under the control of $P_{rpl10}$ (1 kb upstream of the ORF) promoter and $T_{hsp70}$ terminator. |
| pCnVS_NatR$^{FRTm2}$ | 4500 | *Sfi*I-insert of pMF5-1n | Plasmid for expressing dicodon-optimized CnVS with the nourseothricin resistance cassette containing FRTm2 sites (GAAGTTCCTATTCTCAAG AAAGTATAGGAACTTC) at both ends for recycling by Flp recombinase at the *upp3* locus. The expression is under the control of $P_{otef}$ promoter and $T_{nos}$ terminator. |
| pCnVSG_NatR$^{FRTm2}$ | 4356 | *Sfi*I-insert of pMF5-1n | Plasmid for expressing dicodon-optimized CnVS, which is N-terminally fused to eGfp with the nourseothricin resistance cassette containing FRTm2 sites at both ends for recycling by Flp recombinase, at the *upp3* locus. The expression is under the control of $P_{otef}$ promoter and $T_{nos}$ terminator. |
| pCnVSG$^{NES}$_NatR$^{FRTm2}$ | 4499 | *Sfi*I-insert of pMF5-1n | Plasmid for expressing dicodon-optimized CnVS, which is N-terminally fused to eGfp$^{NES}$ with the nourseothricin resistance cassette containing FRTm2 sites at both ends for recycling by Flp recombinase, at the *upp3* locus. The expression is under the control of $P_{otef}$ promoter and $T_{nos}$ terminator. |
| pCop6_NatR$^{FRTm2}$ | 4496 | *Sfi*I-insert of pMF5-1n | Plasmid for expressing dicodon-optimized Cop6 with the nourseothricin resistance cassette containing FRTm2 sites at both ends for recycling by Flp recombinase, at the *upp3* locus. The expression is under the control of $P_{otef}$ promoter and $T_{nos}$ terminator. |
| pCop6G_NatR$^{FRTm2}$ | 4089 | *Sfi*I-insert of pMF5-1n | Plasmid for expressing dicodon-optimized Cop6, which is N-terminally fused to eGfp with the nourseothricin resistance cassette containing FRTm2 sites at both ends for recycling by Flp recombinase, at the *upp3* locus. The expression is under the control of $P_{otef}$ promoter and $T_{nos}$ terminator. |
| pCop6G$^{NES}$_NatR$^{FRTm2}$ | 4316 | *Sfi*I-insert of pMF5-1n | Plasmid for expressing dicodon-optimized Cop6, which is N-terminally fused to eGfp$^{NES}$ with the nourseothricin resistance cassette containing FRTm2 sites at both ends for recycling by Flp recombinase, at the *upp3* locus. The expression is under the control of $P_{otef}$ promoter and $T_{nos}$ terminator. |

### Supplementary Table S4: DNA oligonucleotides used in this study

| Designation | Nucleotide sequence (5'→ 3') | Remarks |
|---|---|---|
| oMB980 | GCAGTCTTGGCGAGCTATTC | *cco1* U1 |
| oMB981 | GGTCTCGCCTGCAATATTTCACCATTATTCTCTTCATTTACTG | *cco1* U2 |
| oMB982 | GGTCTCCAGGCCCGGGCAAATGCCTATCGAG | *cco1* U3 |
| oMB983 | GGTCTCCGGCCGCGAGATGGAAGTGCCTCGC | *cco1* D1 |
| oMB984 | GGTCTCGCTGCAATATTTCTTGCTAGGACTGAAAGCG | *cco1* D2 |
| oMB985 | GACAATGGTGCTTTGCAGGG | *cco1* D3 |
| oMB986 | CTAGGTCTCGTGTCTGAACTCGCACCCAAAGTTG | *cco1* UF_fw |
| oMB987 | CTAGGTCTCAGACACCCTCCGAATGACTATTTGTAC | *cco1* UF_rv |
| oUP054 | CCGCGATTCAGTCAGGTCAG | *cco1* P1 |
| oUP055 | GAATCTATCAGTGACGGCAC | *cco1* P2 |
| oMB996 | GCATCTGTCGCAGCAACATC | *car1* U1 |
| oUP134 | GGTCTCGCCTGCAATATTGATGAATAGCCTCTTTGCCG | *car1* U2 |
| oMB998 | CTAGGTCTCCAGGCCCTTGACCAAGACAAGAACCTCCG | *car1* U3 |
| oMB999 | CTAGGTCTCCGGCCGGAGCGTCTACACAACCGGG | *car1* D1 |
| oUP009 | CTAGGTCTCGCTGCAATAATTTCGACTTCCATACAATGCTCGC | *car1* D2 |
| oUP010 | AAGCTGCTGATGCCGCCTTG | *car1* D3 |
| oUP011 | CTAGGTCTCGTGTCGAGGCGAGGTATAAGCAATG | *car1* DF_fw |
| oUP012 | CTAGGTCTCAGACACCAAGTTGACGTTCTTGTCTC | *car1* DF_rv |
| oUP013 | TCTCCCTGCATGGTAGTAGC | *car2* U1 |
| oUP014 | GGTCTCGCCTGCAATAATACAATAATAAATCGAAGCGGTTCAC | *car2* U2 |
| oUP015 | GGTCTCCAGGCCGACTGGCACTGATTGGTCAAC | *car2* U3 |
| oUP016 | GGTCTCCGGCCAGCATGCACACATGCATCATC | *car2* D1 |
| oUP017 | GGTCTCGCTGCAATATTCACTGCGAAGCGGAGGATC | *car2* D2 |
| oUP018 | CGAAACACAGAAGCGATGAG | *car2* D3 |
| oUP056 | GACAACGCCATGGGACAATC | *car2* P1 |
| oUP057 | CATAGACGGCGTCGACAATG | *car2* P2 |
| oUP186 | CTAGGCGCGCCCCAGCTTTCTACCAAACCGCC | *aat1*_fw |
| oUP187 | CTACCGCGGTTAGTTCTCACGCTTGATAATGATAG | *aat1*_rv |
| oUP266 | CTAGGCGCGCCCTGCAGGATGCTACCTATGTTAC | *hmg1*$^{N\Delta 1\text{-}932}$_fw |
| oUP267 | CTACCGCGGCTATGTGAGAGAAGATGCTCGAC | *hmg1*$^{N\Delta 1\text{-}932}$_rv |
| oUP268 | CTAGGCGCGCCTCGACCGCCACCGTCACCGAG | *idi1*_fw |
| oUP269 | CTACCGCGGTCAGAGGAGGCGGTGAATGC | *idi1*_rv |
| oDD527 | GGTCTCGCCTGCAATATTGCCAATGGCACTAC | *Upa9* U2 |
| oDD605 | GGTCTCGTGGCCAAAGAAGACCAGGCAGCATAG | *Upa9* U3 |
| oDD634 | GGTCTCCGGCCTAGTAGGGTCCAACGTCTAC | *Upa9* D1 |
| oDD635 | GGTCTCCCTGCCAATATTCGACCTCGAGGCCGAG | *Upa9* D2 |
| oMB935 | GATCCCATGGTGAGCAAGGGCGAGGAG | SKL_fw |
| oMB936 | GCGGCCGCTTTAGAGCTTGGACTTGTACAGCTCGTCCAT | SKL_rv |
| oUM121 | CTACCTGCAGGGGTACTGATGACGATGACGAAGAAG | P*rps1*_fw |
| oUM122 | CTACCATGGGATGAATCGATATGTCTTGAGGAAG | P*rps1*_rv |
| oUM123 | CTACCTGCAGGGTATGTGTAGAGGTGGCCTTG | P*rpl10*_fw |
| oUM124 | CTACCATGGCTTGAATACTGTTGGATGGGAGG | P*rpl10*_rv |
| oAB049 | CATGACCAAGAAGTTTGGCACGCTCACCATCT | NES_fw |
| oAB050 | CCGGAGATGGTGAGCGTGCCAAACTTCTTGGT | NES_rv |

### Supplementary Table S5: Codon optimized DNA sequences for expression in *U. maydis*

| **Phytoene synthase *Aa*CrtB from *Agrobacterium aurantiacum*** |
| --- |
| ATGTCGGACCTGGTTCTTACGTCTACCGAAGCTATTACACAAGGCTCGCAGTCGTTCGCTACAGCGGCGAAGCTTATGCCT<br>CCTGGCATCCGCGACGATACAGTCATGCTCTATGCGTGGTGCCGTCATGCTGATGACGTGATTGATGGTCAGGCGCTCGGA<br>AGCCGTCCCGAGGCGGTGAACGACCCACAAGCTCGCCTTGATGGACTCCGCGCAGACACCCTGGCCGCCCTTCAGGGTGA<br>TGGTCCGGTGACCCCTCCATTTGCTGCATTGCGAGCCGTGGCACGACGACACGATTTTCCTCAAGCGTGGCCTATGGATCT<br>CATTGAGGGTTTCGCAATGGATGTTGAGGCTCGTGATTACCGTACACTCGACGACGTGCTGGAGTACAGCTATCATGTTGC<br>CGGAATTGTCGGCGTCATGATGGCCCGAGTCATGGGTGTCCGCGATGATCCTGTGTTGGACCGAGCCTGTGACCTGGGTCT<br>GGCATTCCAGCTGACAAATATCGCACGCGACGTCATCGACGACGCTCGAATTGGACGATGCTACCTTCCAGGAGACTGGC<br>TGGACCAAGCGGGCGCACGAGTGGACGGACCAGTTCCAAGCCCAGAGCTTTACACCGTCATTTTGCGTCTGTTGGACGCA<br>GCCGAATTGTATTACGCCTCTGCACGAGTTGGCTTGGCAGATTTGCCTCCGCGATGCGCGTGGAGCATCGCCGCTGCACTC<br>CGCATCTATCGTGCCATTGGACTGCGTATCCGCAAAGGAGGACCAGAGGCATACCGTCAACGTATTTCCACTTCCAAAGCT<br>GCTAAAATTGGACTCCTGGGCATTGGCGGCTGGGACGTTGCACGCAGCCGCCTCCCCGGCGCGGGTGTCAGCCGCCAAGG<br>CCTGTGGACTCGACCGCACCATGCATAG |
| **(+)-valencene synthase CnVS from *Callitropsis nootkatensis*** |
| ATGGCCGAGATGTTCAACGGCAACTCGAGCAACGACGGCTCGTCGTGTATGCCCGTCAAGGACGCGCTGCGCCGCACCGG<br>CAACCACCACCCCAACCTCTGGACCGACGACTTTATCCAGTCGCTCAACTCGCCCTACTCGGACTCGTCGTACCACAAGCA<br>CCGCGAGATCCTCATCGACGAGATCCGCGACATGTTCTCCAACGGCGAGGGCGACGAGTTCGGTGTGCTCGAGAACATCT<br>GGTTCGTCGACGTCGTCCAGCGTCTCGGCATCGACCGCCACTTCCAGGAGGAGATCAAGACGGCGCTCGACTACATCTACA<br>AGTTCTGGAACCACGATTCGATCTTTGGCGACCTCAACATGGTCGCTCTCGGTTTCCGCATCCTGCGTCTCAACCGCTACGT<br>CGCTTCGAGCGACGTCTTCAAGAAGTTCAAGGGCGAGGAGGGTCAGTTCTCGGGCTTCGAGTCGTCCGACCAGGACGCCA<br>AGCTCGAAATGATGCTCAACCTGTACAAGGCTTCCGAGCTCGACTTCCCCGACGAGGACATTCTCAAGGAGGCGCGTGCTT<br>TTGCTTCGATGTACCTCAAGCACGTCATCAAGGAGTACGGTGACATCCAGGAGAGCAAGAACCCGCTGCTCATGGAGATC<br>GAGTACACCTTCAAGTACCCTTGGCGTTGCCGTCTGCCCCGTCTCGAGGCGTGGAACTTTATCCACATCATGCGTCAGCAG<br>GACTGCAACATCTCGCTCGCCAACAACCTCTACAAGATCCCCAAGATCTACATGAAAAAGATCCTCGAGCTCGCCATCCTC<br>GACTTCAACATCCTCCAGTCGCAGCACCAGCACGAGATGAAGCTCATCTCGACCTGGTGGAAGAACTCGTCGGCCATCCA<br>GCTCGACTTTTTCCGTCACCGCCACATCGAGTCGTACTTCTGGTGGGCCTCGCCCCTCTTCGAGCCCGAGTTCTCGACGTGC<br>CGCATCAACTGCACCAAGCTCTCGACCAAGATGTTCCTGCTCGACGACATCTACGACACCTACGGCACCGTCGAAGAGCTC<br>AAGCCCTTCACCACCACGCTCACCCGCTGGGACGTCTCGACCGTCGACAACCACCCCGACTACATGAAGATCGCCTTCAAC<br>TTTAGCTACGAGATCTACAAGGAGATCGCCTCGGAAGCTGAGCGCAAGCACGGTCCTTTTGTCTACAAGTACCTCCAGTCG<br>TGCTGGAAGTCGTACATCGAGGCGTACATGCAGGAGGCCGAGTGGATCGCCAGCAACCACATCCCTGGTTTCGACGAGTA<br>CCTCATGAACGGTGTCAAGTCGTCGGGTATGCGCATCCTCATGATTCACGCGCTCATCCTCATGGACACGCCGCTCTCCGA<br>CGAGATTCTCGAACAGCTCGACATCCCTTCGTCGAAGTCGCAGGCTCTGCTCTCGCTCATCACGCGTCTCGTCGACGACGT<br>CAAGGACTTTGAAGACGAGCAGGCGCATGGCGAGATGGCTTCGTCGATCGAGTGCTACATGAAGGACAACCACGGTTCGA<br>CGCGCGAGGATGCGCTCAACTACCTCAAGATCCGCATCGAGTCGTGCGTCCAGGAGCTCAACAAGGAGTTGCTCGAGCCC<br>TCGAACATGCACGGCTCGTTCCGCAACCTCTACCTCAACGTCGGCATGCGTGTCATCTTCTTCATGCTCAACGACGGTGAC<br>CTCTTCACCCACTCGAACCGCAAGGAGATCCAGGACGCCATCACCAAGTTCTTTGTCGAGCCCATCATCCCCTGA |
| **The fungal sesquiterpenoid synthase Cop6 from *Coprinopsis cinerea*** |
| ATGCCTGCTGCTCTGCCCTACAACGTCTCGCGCGACAACAAGTGGGATATCAAGAAGATCATCCAGGACTTTTTCAAGCGC<br>TGCGATGTGCCCTACCAGGTCATCCCCTACGACACCGAGCTCTGGAACGCCTGCCTCAAGCGTGCCAAGGAGAAGGGCTA<br>CCCCGTCGAGCCCGACTCGCCCATGTCGCTCTACCGCAGCTTCAAGGTCGGTGTCGTCATCACGCGCACCTCGTACGGTCA<br>CATCCAGGACTACGAGATCCTCATCTGGGTCGCCACCTTCACCGCCTTTGTCACCTACGCCGACGACGCCTTCCAGGAGGA<br>CATCCAGCACCTCCACAGCTTCGCTCGCACCTTCCTCCAGAACGAAAAGCACGAGCATCCCGTCCTCGAGGCCTTTGCCCA<br>GTTCCTGCGCGAGTCGTCGATCCGATTCTCGCACTTTGTCGCCAACACCGTCGTCTCGTCGGCGCTGCGCTTCATGATGTCG<br>ATCGCGCTCGAGTTCGAGGGTCAGAACGTCTCGGTCTCGACCGAGGCGCGTGAGTACCCTGGCTACATCCGCATCCTCTCG<br>GGTCTCTCGGACATCTACGCGCTCTTCGCCTTCCCCATGGACCTGCCGCGTTCCACCTACATCCAGGCCTTCCCCGAGCAGA<br>TCGACTACATCAACGGCACCAACGACCTGCTCTCGTTCTACAAGGAGGAGCTCGACTGCGAGACCGTCAACTTTATCTCGG<br>CCGCTGCCACCTCGCAGCAGGTCAGCAAGCTCGAGGTGCTGCGCAACGCCGCCGAGAAGGCCGCCTACTCGTACGACGTC<br>GTCGTCAACGTGCTCAAGCCCTACCCCGAGGCGCTTGCCGCCTGGAAGTCGTTTGCTCGCGGCTTCTGCTACTTCCACACCT<br>CGTCGCCACGCTACCGTCTCGGCGAGATGTTCCACGACTTTGAGCACGACCTCGTCTGCAAGTGCGCCTCGTGCACCGAGA<br>TCTGA |
